# Timing Is Everything In Autumn – Coordination Of Senescence Onset By A Transcriptional Program In Response To Environmental And Phytohormone Signals

**DOI:** 10.1101/2022.03.30.486406

**Authors:** Jenna Lihavainen, Jan Šimura, Pushan Bag, Nazeer Fataftah, Kathryn Megan Robinson, Nicolas Delhomme, Ondřej Novák, Karin Ljung, Stefan Jansson

## Abstract

European aspen (*Populus tremula* L.) undergoes a coordinated senescence program during autumn; however, it is not known what exactly triggers it. To identify the cellular program leading to senescence, we utilized natural variation among Swedish aspen genotypes in a common garden to study senescence timing and the underlying changes in leaf phytohormone and transcriptome profiles. Apart from the patterns of major transcriptional cascade that was similar between the genotypes and closely associated with cytokinin and auxin metabolite levels and gradually decreasing air temperature during autumn, we detected patterns that consistently preceded or coincided with senescence onset in individual genotypes. Another cascade seemed to respond to short-term changes in weather conditions that re-wired the transcriptional network; the up-regulation of genes related to ethylene and abiotic stress, programmed cell death and translation occurred first in the early-senescing genotypes and later in the late one. Network analyses displayed a connection between the two cascades, metabolic stress and immunity responses mediated by salicylic acid (SA)-signalling pathway that was repressed along with SA levels at senescence onset. We propose that autumn senescence in aspen trees is affected by environmental variation that evokes stress and the timing is fine-tuned by their stress tolerance mechanism.

**One sentence summary:** Salicylic acid signalling pathway fine-tunes the timing of senescence onset under challenging environmental conditions in autumn.

## INTRODUCTION

Deciduous trees in temperate regions salvage nutrients through senescence before abscission of the leaves in autumn, which has a massive ecological importance. Although beautiful autumn colours are widely appreciated by the public, senescence regulation at the molecular level is not well understood. Research efforts (reviewed in Gepstein et al., 2004; Lim et al., 2007; Woo et al., 2013), such as genetic approaches (to identify genes where mutations lead to premature or delayed senescence), transcriptomics (to identify senescence-associated genes - SAGs), external applications of phytohormones, and profiling of metabolite levels and reactive oxygen species (ROS) have connected them with senescence (Balazadeh et al., 2008; Buchanan-Wollaston et al., 2003; Buchanan-Wollaston et al., 2005; Watanabe et al., 2018; Zhang and Guo, 2018). Yet, there is no consensus on the senescence trigger, which metabolites, genes or post-translational mechanisms are the most important ones – across species – for the process, or which environmental factors could consistently explain why senescence starts at a certain time in trees.

In our previous studies of autumn senescence in European aspen (*Populus tremula* L.) we found that a given aspen genotype in its natural environment typically initiates senescence (determined by rapid chlorophyll depletion) on around the same date every year (Keskitalo et al., 2005; Fracheboud et al., 2009). However, the date shifts if the same genotype is instead grown either in a greenhouse (Fracheboud et al., 2009), or at another latitude (Michelson et al., 2018), or if the tree is girdled (Lihavainen et al., 2020) or fertilized (Fataftah et al., bioRxiv). Moreover, with multi-year studies of one genotype, we previously proposed that 1) the overall transcriptome profile can be difficult to link with senescence onset (Jansson and Thomas, 2008) and 2) that cytokinin levels may not explain senescence onset under natural conditions (Edlund et al., 2017). These observations suggest that senescence in deciduous trees under natural conditions cannot be explained by either global transcriptome patterns or by a single hormone signaling pathway; rather it is most likely affected by an intricate regulatory network that integrates external environmental cues and internal metabolic signals.

On one hand, a problem with our approach of studying senescence in natural conditions may be that in nature many factors vary at the same time, which makes it difficult to separate the “true signal” from the environmental variation. On the other hand, our approach can have several benefits compared to experiments performed under controlled conditions, as the latter cannot separate a developmental program from a timetable (Jansson and Thomas, 2008), that mature trees – that cannot be easily grown under controlled conditions – have more “stable” senescence patterns and are likely to integrate multiple signals to know that it is autumn. Therefore, by taking advantage of our previous study of autumn phenology in populations of natural aspen accessions sampled from a large latitudinal range (Fracheboud et al., 2009, Michelson et al., 2018), we aimed to understand the complex regulatory program leading to senescence onset in several genetically different aspen trees that vary substantially in their senescence onset dates in a common garden. Theoretically, the changes that truly regulate the onset of autumn senescence should precede or coincide with the onset, first in the genotype that starts to senesce first and last in the genotype that senesces last, assuming that they utilize similar transcriptional or hormonal programs to integrate the signals to initiate senescence.

Here, we have used the most recent gene models in the aspen genome and co-expression networks to study phytohormone and transcriptome changes during autumn in several aspen genotypes. We demonstrate that there are two major transcriptional cascades changing in parallel in autumn, and they respond to environmental variation and phytohormones. Both are interlinked with the salicylic acid (SA)-signaling pathway that appears to play a crucial role in the regulation of senescence onset in aspen under stressful environmental conditions in autumn through its connection with other phytohormones and post-transcriptional stress tolerance mechanisms.

## MATERIAL AND METHODS

### Plant material

European aspen (*Populus tremula*) genotypes selected for this study were genotype 201, a mature tree situated at Umeå University campus, and five other genotypes belonging to the Swedish aspen collection (SwAsp, planted 2004, Luquez et al., 2008). Clonal replicates of 201 are a part of Umeå aspen collection (UmAsp, planted 2009-2010) and like SwAsp trees they were grown in common garden at the field station of Forest Research Institute in Sävar (63.9°N) near Umeå. SwAsp genotypes (1, 33, 48, 81 and 96) originating from different latitudes were selected for the study based on their senescence phenotypes. Each SwAsp genotype had at least three individual trees growing in the field, and they were 14 years old at the time of sampling in autumn 2018.

### Weather parameters and solar spectral quality

Weather data (air temperature, solar radiation, precipitation, air pressure, and relative humidity) for study years 2011 and 2018 were obtained from the Swedish Meteorological and Hydrological Institute (SMHI, http://shmi.se) and from the TFE weather station (http://www8.tfe.umu.se/TFE-vader/, Department of Applied Physics and Electronics, Umeå University). Solar irradiance and spectral quality were measured with an ILT900-R Spectroradiometer (InternationalLight Technologies) on 14 dates during autumn 2020 between 240-268 DOY (the day of the year). Cloud cover data were based on an oktas scale, where 0 represents a clear sky and 8 is fully overcast (SHMI). Measurements were performed on sunny days with a clear or almost clear sky (scale 0-2, n=3), partly cloudy days (scale 3-5, n=6), and on almost fully overcast cloudy days (scale 6-8, n=5) at Umeå University campus. Light spectra were measured 3-6 times around noon (12 am-1 pm). Each recorded measurement at 1 nm intervals was computed as an average of ten scans. Wavelength bands were assigned as blue (450-495 nm), red (620-700 nm) and far-red (700-750 nm). Those wavelength bands were used to calculate blue-to-red (B:R) and red-to-far-red (R:FR) ratios for sunny, partly cloudy and overcast days. Saturated vapour pressure (SVP) at given temperature (*T*, hourly mean) was calculated based on the equation: SVP=(0.6108*exp[*T*\*17.27/*T*+237.3]) and air vapour pressure deficit (VPD) was derived from the equation: VPD=SVP*(1-RH/100), where RH is hourly mean relative air humidity (%).

### Sampling

All aspen collections are grown in a common garden in Sävar, approx. 15 km distance from Umeå. Leaves were sampled twice a week from 6^th^ of August (218 DOY) until 27^th^ of September (270 DOY) in 2018. In genotype 96, leaf sampling was omitted on 270 DOY, since most of the leaves had abscised. Samples were collected from three individual trees per SwAsp genotype (n=3). Sampling was always performed at noon and finished within one hour (12 am - 1 pm). Five healthy-looking short-shoot leaves were sampled from the middle level of the canopy (from four sides of the tree) using a telescopic tree pruner. The leaves were pooled, wrapped in aluminium foil and frozen immediately in liquid nitrogen. The sampling of leaves from genotype 201 situated at Umeå University campus in autumn 2011 was similar, as described in Edlund et al., (2017).

### Chlorophyll content and the onset and rate of autumn senescence

Chlorophyll content index (CCI) of leaves (mean of five leaves) of the five SwAsp genotypes was determined twice a week from 6^th^ of August (218 DOY) until 9^th^ of October (282 DOY) in autumn 2018 with a chlorophyll meter (CCM 200 plus, Opti-Sciences). The CCI data of SwAsp genotypes in the field in 2011 was obtained from Michelson et al., (2018) and in the greenhouse study in 2006 from Fracheboud et al., (2009). The CCI data of genotype 201 in 2011 was obtained from Edlund et al., (2017) and the CCI values (mean of five leaves) of its clonal replicates grown in common garden were recorded in autumn 2018. In all cases, the senescence onset dates in autumn were determined based on the chlorophyll content index (CCI) as the day of the year (DOY) when rapid chlorophyll depletion started, using the curve fitting method described in Lihavainen et al., (2020).

### RNA extraction and mRNA sequencing

The leaves were ground to fine powder in liquid nitrogen with a mortar and a pestle. Total RNA was extracted from 500 mg of leaf sample from three SwAsp genotypes (1, 48 and 81) from ten time points during autumn (225-270 DOY) 2018 using Spectrum™ Plant Total RNA isolation kit (Sigma-Aldrich) according to manufacturer’s protocol (n=3 individual trees per genotype and time point). DNA was removed with DNA-*free*™ DNA removal kit (Thermo Scientific). All the RNA samples were of good quality with OD 280/260 ≥2.0 assessed with 2000 Nanodrop (NanoDrop Technologies, Wilmington, DE, USA) and RNA integrity number (RIN) ≥8.0 determined with Agilent 2100 BioAnalyzer (Agilent Technologies, Waldbronn, Germany). RNA concentration was determined with Qubit fluorometer 2.0 using Qubit™ RNA BR Assay Kit (Thermo Scientific). mRNA sequencing was conducted by Science for Life Laboratory in Stockholm, Sweden (SciLifeLab) using the Illumina NovaSeq 6000 platform. Raw data can be retrieved from the European Nucleotide Archive (https://www.ebi.ac.uk/ena/browser/home) under the accession PRJEB51801.

### Transcriptomic data processing

The data pre-processing steps were performed following the method based on Delhomme et al., (2014). The steps included quality control where the raw sequence data was first assessed (FastQC v0.10.1), and ribosomal RNA was removed (SortMeRNA v2.1b) (Kopylova et al., 2012) followed by trimming in order to remove adapter sequences (Trimmomatic v0.32) (Bolger et al., 2014) after which another quality control step was performed to ensure that technical artefacts were not introduced during the pre-processing steps. mRNA data were aligned with Salmon (v0.14.2) (Patro et al., 2017) using *Populus tremula* v2.2 cDNA library as a reference. Principal Component Analysis (PCA) and hierarchical clustering analysis (HCA) were applied in R (version 4.0.0, R-Core-Team 2015) to assess the similarity among the biological replicates (biological quality control). One outlier sample had low library size and based on the PCA, samples from 270 DOY that were from yellowing leaves, were separated from the other samples. In order to reduce their contribution to the overall variation, they were excluded from the following analyses along with the outlier sample. For visualisation and downstream analyses, the counts were normalized using the variance stabilizing transformation (VST) implemented in the Bioconductor DESeq2 package (v1.16.1, Love et al., 2014). We have previously performed mRNA sequencing with one aspen tree (I201) growing in Umeå University campus (Edlund et al., 2017, Gene Expression Omnibus [GEO] accession number GSE86960) and here, the RNA sequencing data was re-aligned with Salmon as described above to compare the data sets from two study years. An overview of the mRNA data processing steps and parameters are available in the GitHub repository (https://doi.org/10.5281/zenodo.5906743).

### Global gene expression patterns during autumn

The overall variation in transcriptome data in autumns 2018 and 2011 was visualised using principal component analysis (PCA, SIMCA P+, version 15, Umetrics, Umeå, Sweden). Data were log_10_-transformed and scaled by unit variance. In addition to scores plots, the changes in global gene expression profiles were visualised as time-dependent plots of PC scores.

### Weighed gene co-expression network analysis

Weighed gene co-expression network analysis (WGCNA) was performed to identify gene modules and hub genes that are candidates to regulate biological processes in aspen leaves in autumn using the WGCNA package in R (Langfelder and Horvath, 2008, Langfelder et al., 2011). See details in Supplemental Methods.

In short, WGCNA was conducted with 21 602 genes that displayed temporal variation and were expressed in all genotypes in 2018, and with 19 287 genes in 2011. Differential analysis was performed to study the preservation of the gene interconnectivity among the three genotypes in 2018. To study relationships among gene expression patterns, environmental factors and metabolic markers, weather parameters, chlorophyll content index (CCI), and phytohormone levels were integrated in WGCNA and their correlation with eigengene modules as well as individual genes in each genotype and in the consensus network were interpreted. Hub genes with high intramodular connectivity were identified based on network topology structure. The relationships between eigengene modules and their correlations with weather parameters and chlorophyll levels were confirmed with similar WGCNA analysis performed with another genotype in another year.

Gene Ontology (GO) term enrichment analyses were performed for genes in each eigengene module with PlantGenIE using the aspen database (https://plantgenie.org, Sundell et al., 2015). Since a greater proportion of Arabidopsis genes have annotation than do aspen genes, complementary GO term enrichment analyses and PFAM (protein family and domain) we performed using the best DIAMOND hits for Arabidopsis genes (PlantGenIE). Kyoto Encyclopedia of Genes and Genomes (KEGG) pathway enrichment analysis was performed with g:Profiler (https://biit.cs.ut.ee/gprofiler/gost, Raudvere et al., 2019) and GO term networks were produced with ClueGO (version 2.5.7, Bindea et al., 2009) and CluePedia (version 1.5.7, Bindea et al., 2013) applications in Cytoscape (version 3.8.0) using the best DIAMOND hits for Arabidopsis gene IDs (Shannon et al., 2003). Co-expression networks were produced for each SwAsp genotype separately that were then merged in Cytoscape to visualise consistent relationships (edges present in all three genotypes with FDR-adjusted P-value <0.05). For clear network visualisation, edges with absolute correlation coefficient > 0.6 were included to focus on the connections between close neighbours. Co-expression networks were visualised so that the size of the node is proportional to its degree (connectivity), with larger size depicting higher degree. Nodes with high degree are considered as important hubs with potential regulatory roles and nodes with high betweenness centrality as important mediators controlling the information flow through the network structure.

### Differentially expressed genes

Differentially expressed (DE) genes across time points and between genotypes in autumn 2018 were determined with the DESeq2 package, with false discovery rate (FDR) adjusted *P*-value <0.01 considered significant in all cases. Time effect between specific time points was studied in two ways; by comparing time points against the first time point (225 DOY) and by comparing consecutive time points considering the whole data (all genotypes) set and within each genotype (log_2_ fold change [FC] cut off >0.5, Schurch et al., 2016). The difference in gene expression levels between individual genotype pairs was also tested. In addition, the main effects of time (between any time points), genotype (between any genotype) and their interaction effect (time×genotype *i*.*e*., different temporal patterns among the genotypes) were studied using likelihood ratio tests (LRT).

Senescence-associated genes (SAGs) were identified by comparing the first and the last time points. Since the genes showed considerable variation in their expression between consecutive time points (>0.5 log_2_FC), gene lists were filtered based on cut-off >1.0 to find genes with a consistent increase or decrease over the autumn. The intersecting set of DE genes was visualized using Venn diagrams (Heberle et al., 2015). GO term enrichment analysis was performed for identified DE gene groups as described above.

### Meta-analysis of senescence associated genes in Populus spp. during autumn

A meta-analysis was performed to study the overlap between up- and down-regulated genes during autumn in aspen (*Populus tremula*) in 2011 (genotype 201) and 2018 (genotypes 1, 48 and 81) and in *Populus* spp. Genes that showed log_2_FC >1.0 between the first and the last time points (217-265, 225-264 DOY) in aspen were regarded as senescence associated genes (SAGs). The lists of consistent up- and down-regulated genes in all aspen genotypes were compared with genes differentially expressed in poplar (*P. trichocarpa*) leaves during autumn (Li et al., 2020, Lu et al., 2020). The lists of up- and down-regulated genes in poplar were obtained from the Leaf Senescence Database (Li et al., 2020, LSD 3.0, https://bigd.big.ac.cn/lsd/poplar.php) and from Lu et al., 2020. For comparative analyses, *Populus tremula* gene IDs (Potra gene ID) were converted into poplar gene IDs (Potri gene ID) using the best DIAMOND hits obtained from PlantGenIE. The number of unique and common up- and down-regulated genes during autumn between the two study years and between the two *Populus* species were visualized using Venn diagrams. GO term enrichment and network analyses were performed for identified gene sets as described above.

### Phytohormones

Metabolites were extracted from samples collected in autumn 2018 from five SwAsp genotypes (n=2-3 individual trees per genotype and time point). Leaf samples (∼10 mg FW) were extracted with 1 mL of ice cold 10% MeOH with 1 M formic acid (FA). Labelled internal standards (10 µL) and two tungsten beads (3 mm diameter) were added to each sample and samples were homogenized for 10 min in a bead mill (25 Hz, Retsch Qiagen). Samples were centrifuged for 15 min at 14 000 rpm at 4°C. The supernatant was divided in two 450 µL aliquots that were diluted to 1 mL with 1 M formic acid (FA) and with water for analysis of cytokinins (CKs) and stress induced phytohormones (SIP, jasmonic acid, auxin, salicylic acid, abscisic acid and metabolites), respectively. The purification of isoprenoid CKs was carried out according to (Dobrev and Kaminek, 2002) using the Oasis MCX column (30 mg of C18/SCX combined sorbent with cation-exchange properties, Waters Inc., Milford, MA, USA). Analytes were eluted by two-step elution using a 0.35 M NH_4_OH aqueous solution and 0.35 M NH_4_OH in 60% (v/v) MeOH solution. and the other set of samples for SIP analysis was purified using the Oasis HLB (30 mg 1 cc, Waters Inc., Milford, MA, USA) as described in Floková et al., (2014). The eluates were collected in test tubes, evaporated to dryness using speed vac (SpeedVac SPD111V, Thermo Scientific, Waltham, MA, USA) and stored at -70°C.

Prior to LC-MS analysis, samples were dissolved in 40 µL of 30%MeOH and analysed with the UHPLC-ESI-MS/MS system comprising of a 1290 Infinity Binary LC System coupled to a 6490 Triple Quad LC/MS System with Jet Stream and Dual Ion Funnel technologies (Agilent Technologies, Santa Clara, CA, USA). The UHPLC-ESI-MS/MS parameters used to analyse cytokinin and stress induced phytohormone levels were described in Svačinová et al., 2012 and Floková et al., 2014, respectively. Phytohormone levels were quantified by normalizing analyte peak area by labelled internal standard peak area, using a calibration curve of the reference standard and the fresh weight of the sample using Agilent MassHunter Workstation Software Quantitative (Agilent Technologies, Santa Clara, CA, USA). The concentrations of detected 23 phytohormones were expressed as pmol g^-1^ FW.

The effects of genotype, time (DOY) and their interaction on phytohormone levels in autumn 2018 were tested with log_10_-transformed data with two-way ANOVA, *P*-value <0.05 considered significant (IBM SPSS Statistics version 25).

## RESULTS

### Early, intermediate- and- and late-senescing genotypes were consistently different in the field and in the greenhouse

We have extensively studied the variation in autumn phenology and senescence onset dates in genotypes from the Swedish aspen collection (SwAsp) in a common garden and in a genotype I201 growing at the Umeå University campus (part of the Umeå aspen collection - UmAsp) (Keskitalo et al., 2005, Luquez et al., 2008, Fracheboud et al., 2009, Michelson et al., 2017, Wang et al., 2018). Therefore, for this study, we selected the genotype I201 and five SwAsp genotypes originating from southern (L1, L33), middle (I48) and northern (E81, E96) latitudes of Sweden that showed high variation in senescence onset dates (Fig. 1A). In 2011, I201 started to senescence around 255 DOY within the typical range for this genotype (Fig. 1, Keskitalo et al., 2005, Fracheboud et al., 2009) and in 2018, the clonal replicates of I201 in the common garden showed similar onset dates to the parent tree (Fig. 1C, Fig. S1) confirming that that onset of senescence was independent of tree age. In autumn 2018, genotype E81 started to senescence around 242 DOY, E96 on 244 DOY, I48 on 247 DOY, L33 on 263 DOY and L1 on 264 DOY (Fig. 1B) close to the mean senescence date over the two study years (2011 and 2018, Fig. 1). Based on the senescence timing in both aspen collections, these genotypes were classified as early-(around or before 245 DOY, E81, E96), intermediate-(between 245-260 DOY, I48 and I201) and late-senescence (after 260 DOY, L1, L33) phenotypes (Fig. 1). The onset of senescence in a greenhouse under natural photoperiod and light regime in the three genotypes (E81, I48, L1) selected for the transcriptome profiling showed similar relative variation as in the field, showing that it was stable also under constant temperature conditions (Fig. S2).

**Figure 1.**
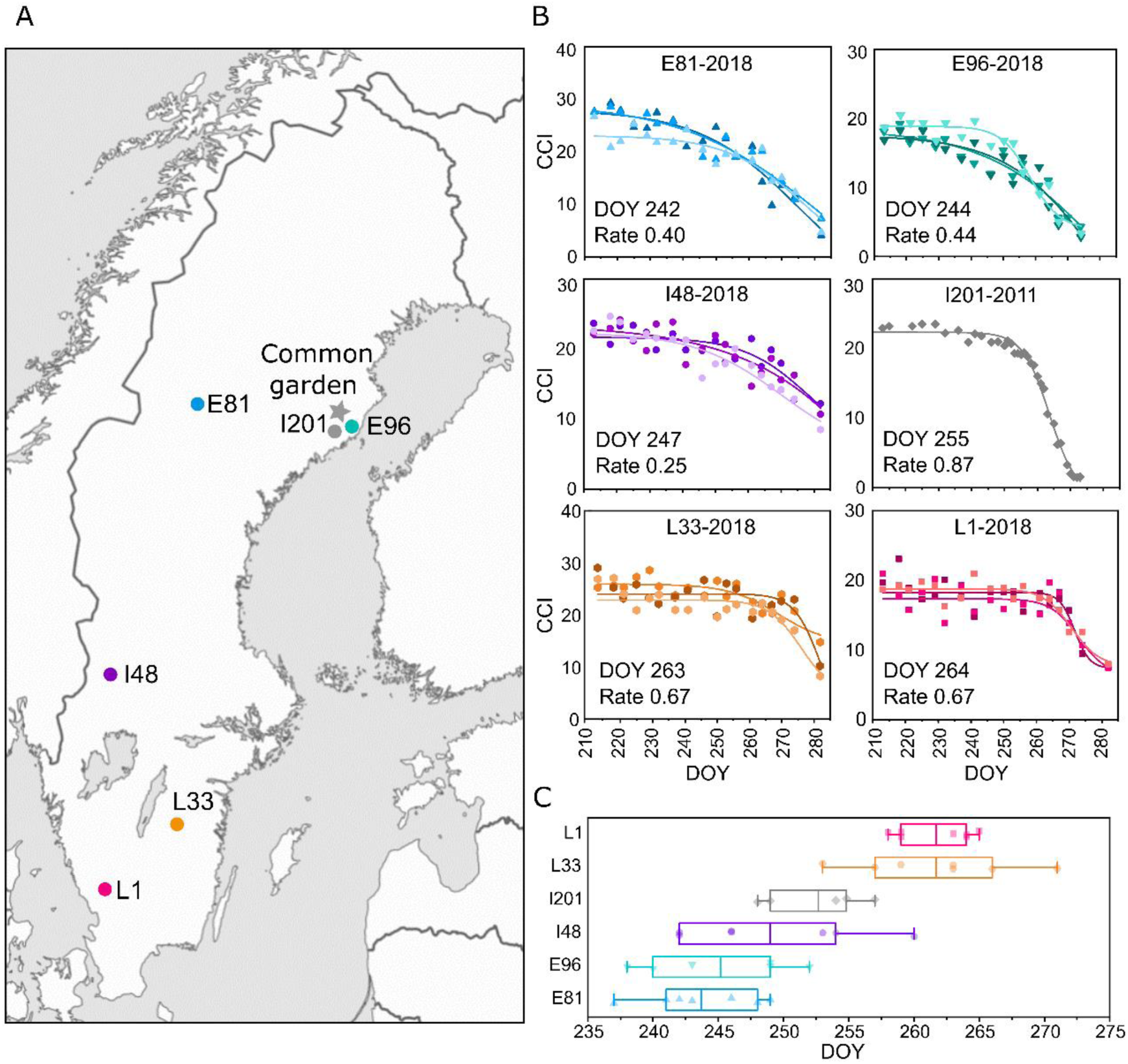
Autumn senescence phenotypes. Swedish aspen (SwAsp) genotypes originating from the south (L1, L33), central (I48) and north (E81, E96) of Sweden were grown in a common garden in Sävar near Umeå (marked with a star, A). Genotype I201 is local to Umeå, situated at Umeå University campus, and its clones are part of Umeå aspen collection grown in a common garden in Sävar (A). The onset and the rate of autumn senescence were determined based on the chlorophyll content index (CCI). Chlorophyll curves (CCI values as a mean of five leaves) are shown for the campus tree, I201, in autumn 2011 and for the three replicate trees of SwAsp genotypes in autumn 2018 (B). The variation in senescence onset date over the two study years are shown in a box plot representing mean (solid line), 25% and 75% quartiles and minimum and maximum values (whiskers) (C). Dots present data over two study years, data are from one parent tree (2011) and four clonal replicates in UmAsp collection (2018) of genotype I201 (n=5), and from three to four replicates of SwAsp genotypes in each study year (n=6-7).

### Neither major variation in the transcriptome profile nor senescence-associated gene expression explain senescence onset

Transcriptome profiles of one each early-(E81), intermediate-(I48) and late-senescence genotypes (L1) in autumn 2018 were compared using differential expression (DE) analyses (Fig. 2), multivariate statistics (Principal Component Analysis, PCA, Fig. 2, Figs. S3 and S4) and weighed gene co-expression network analysis (WGCNA details in Supplemental Methods, result details in Supplemental Data Set 1 and 2 and Figs. S5-14) that unlike PCA considers the variation and gene relationships in individual genotypes as well as in a consensus network. PCA and WGCNA were performed separately with the transcriptome data from genotype I201 in autumn 2011 and the data from the two study years were then compared to find consistent patterns.

**Figure 2.**
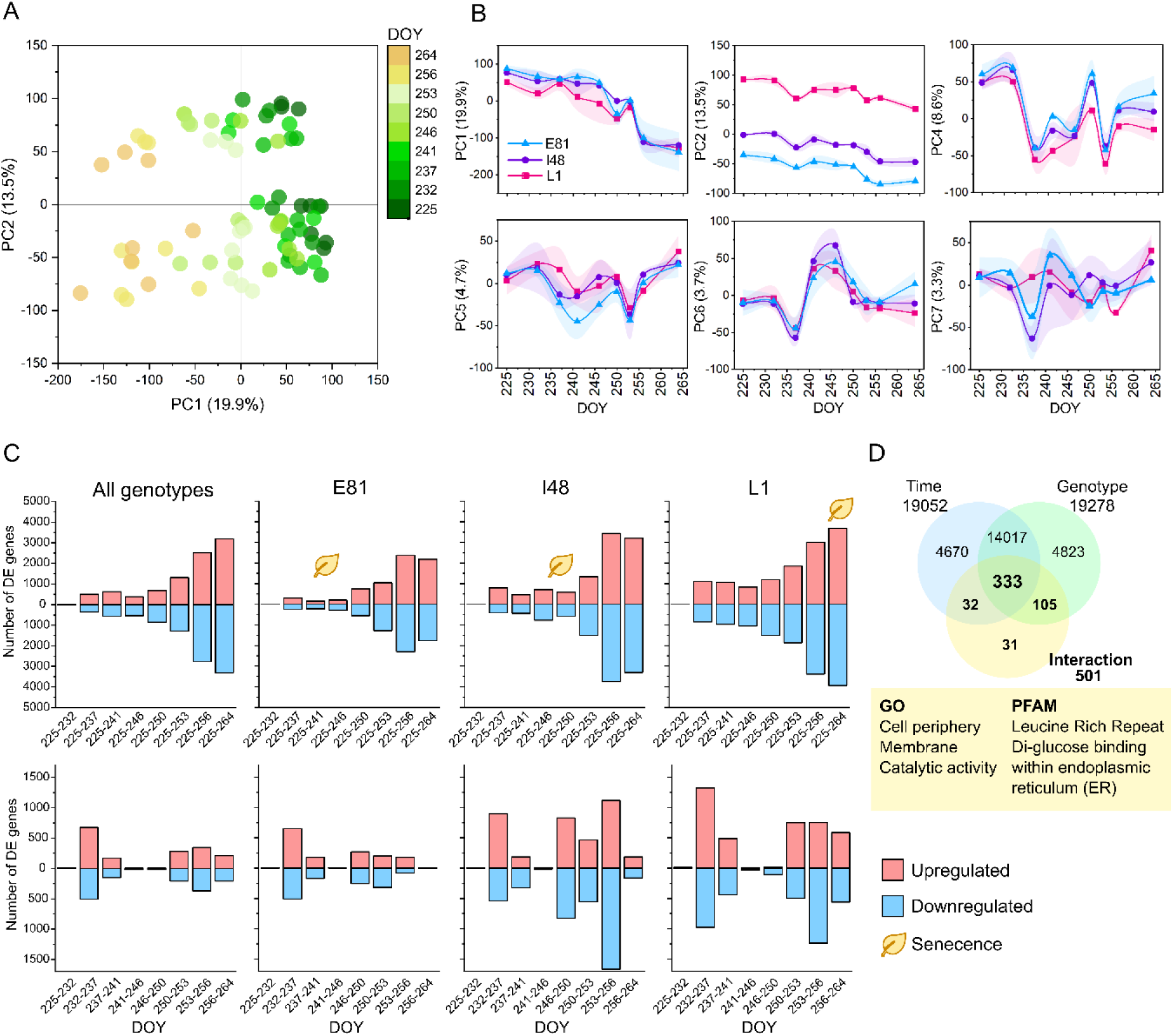
Global transcriptome profile and differentially expressed (DE) genes in aspen leaves during autumn 2018. Principal component analysis (PCA) of the transcriptome profile in three SwAsp genotypes in 2018 (A) and time-dependent patterns of principal component (PC) scores (B). Data are mean ± SE (shadowed area), n=2-3. The effect compared to the first time point (225 DOY) and between consecutive time points on gene expression levels were tested with DESeq2 (C). Time effect was tested considering the whole data with all genotypes (consensus) and within each genotype (FDR adjusted *P*<0.01, log_2_ fold change cut off >0.5) (C).. Venn diagram depicts the number and overlap of genes with significant time, genotype and interaction effect based on likelihood ratio test (LRT, D), FDR adjusted *P*<0.01 considered significant. Enriched Gene Ontology (GO) terms and PFAM (protein families and domains) for the gene set (501) with significant interaction effect (D). The time-dependent score plots of all nine significant PCs are in Fig. S3, the list of DE genes and the details of enrichment analyses are in Supplemental Data Set 4.

Pronounced temporal patterns in the transcriptome were – as expected – found in all genotypes and both study years. PCA yielded two and nine principal components (PCs) and WGCNA analysis 23 and 27 eigengene modules with distinct genotype-or time-related patterns in 2011 and 2018, respectively (Figs. 2 and 3, Figs. S3, S11, S13). In both study years the major principal components (the first PCs) that explain most of the variation in the data displayed gradual shifts in the global transcriptome profiles (Fig. 2, Fig. S4). If this shift was responsible for the timing of senescence by initiating it, it would have appeared earlier in the early-senescing genotype E81 than in the late-senescing L1 reflecting the difference of 23 days in their senescence onset dates. However, instead, the shift in the gene expression profile and the increase in the number of DE genes occurred earlier in L1 than in the other two SwAsp genotypes (Fig. 2) indicating that the main variation in global transcriptome profile did not explain senescence onset.

**Figure 3.**
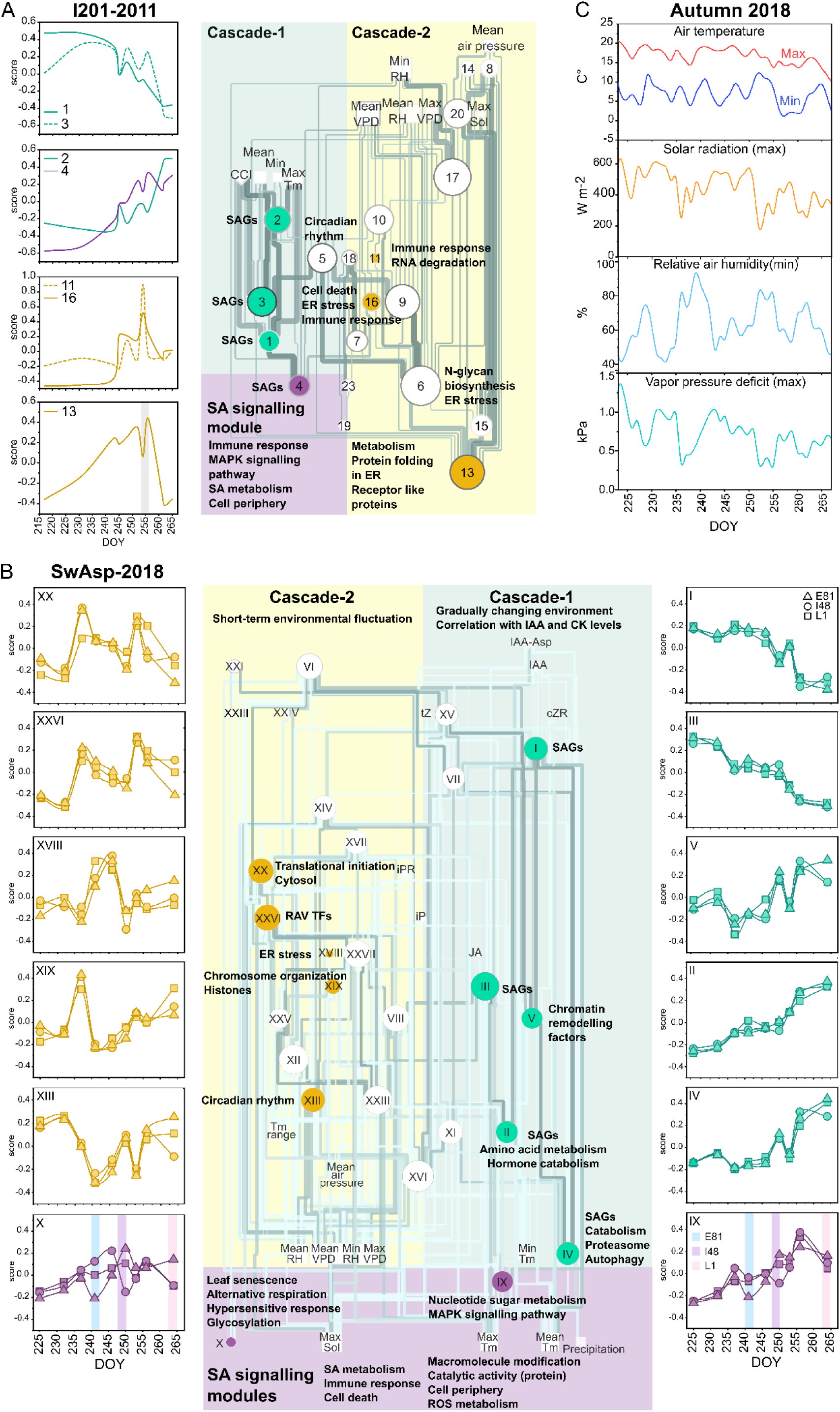
Transcriptome patterns and their relationship with weather parameters and phytohormone levels during autumn. Weighted gene co-expression network (WGCNA) of eigengene modules, weather parameters (past 24 h) and phytohormone levels in autumn 2011 (A) and 2018 (B) and the temporal patterns of selected modules. Module expression patterns in genotype I201 in 2011 are from one tree (n=1) and in 2018, the patterns in three SwAsp genotypes are from 2-3 trees (B, mean of n=2-3 in each time point per genotype). Coloured line represents senescence onset in a particular genotype. Enriched Gene Ontology (GO) terms and Kyoto Encyclopedia of Genes and Genomes (KEGG) pathways are shown beside the nodes. The significant relationships (edges with *P*-value <0.05) are shown and coloured based on the correlation coefficient (darker colour depicts higher absolute Pearson r). In SwAsp network, the edges common for all three genotypes are shown (B). The size of the node is proportional to its degree, larger node size depicting higher connectivity. Modules are numbered based on the descending number of assigned genes. Minimum and maximum air temperature (°C), maximum solar radiation (W m^-2^), minimum relative humidity (RH%) and maximum air vapour pressure deficit (VPD) during the study period in autumn 2018 (C). All weather parameters, phytohormone levels and eigengene patterns along with the network structures displaying positive and negative relationships are in Figures S7, S8, S11 and S13. The abundances of transcription factors and chromatin remodelling factors and enrichment analyses are in Supplemental Data Set 1 (2018) and 2 (2011). ER =endoplasmic reticulum, UPR =unfolded protein response, SAGs=senescence-associated genes, SA=salicylic acid, CK=cytokinins (Z=zeatin, iP=isopentenyl adenine, DHZ=dehydrozeatin), IAA=auxin, JA=jasmonic acid

To identify the underlying sources of variation in the transcriptome, we investigated WGCNA eigengene module and gene expression relationship with weather parameters and chlorophyll levels in the two study years (Fig. 3, Figs. S9-S10, Figs. S14). Network structures of both years displayed broadly two major cascades changing in parallel in autumn (Fig. 3AB). In both years, the first one (cascade-1) involved the largest gene modules with gradually enhanced or repressed expression during autumn (Fig. 3AB). In autumn 2018, a module enriched with genes encoding chromatin remodelling factors was associated with this cascade (Fig. 3B). The genes in the up-regulated modules were associated with protein trafficking and catabolism, autophagy, phytohormone metabolism and stress responses (Fig. 3AB, Fig. S11, Fig. S13, Supplemental Data Set 1 and 2). The hub genes in the modules in the SwAsp genotypes encoded typical senescence-associated genes (SAGs) such as *WRKY75, NYE1 (NON-YELLOWING1)*, and *ATAF1* (Supplemental Data Set 1). Down-regulated gene modules contained genes involved in photosynthesis, chloroplastic functions and major biosynthetic pathways (Fig. 3, Fig. S11, Fig. S13, Supplemental Data Set 1 and 2). Those modules followed similar gradually changing patterns correlating with decreasing air temperature in all genotypes (Fig. 3AB). Those patterns followed chlorophyll levels only in the genotypes that senesced in early (E81) or mid-autumn (I48, I201), but not in the genotype that senesced in late autumn (L1, Fig. S15). Therefore, they could not explain the different timing of senescence onset in a consistent manner.

Next, we looked for senescence-associated genes (SAGs) that were up- or down-regulated during autumn, and that could potentially trigger senescence without affecting overall transcriptome profiles, but by re-wiring the regulatory network. In our dataset, the expression of 511 and 353 SAGs were consistently enhanced or repressed during autumn, respectively, over the two years (Fig. 4A, Supplemental Data Set 3). Those SAGs were consistently assigned into the modules in cascade-1 in both years. As our PlantGenIE database makes it possible, with good precision, to identify homologous genes in the aspen and poplar (*P. trichocarpa*) genomes, we compared our results with those obtained from two studies of poplar (Leaf Senescence Database, Li et al., 2020, LSD 3.0, https://bigd.big.ac.cn/lsd/poplar.php; Lu et al., 2020, Fig. 4B). SAGs with enhanced expression (95 genes) during autumn in both species were related to innate immunity encoding WRKY, NAC and TGA TFs and the repressed genes (121) were enriched with GO terms related to chloroplastic processes (Fig. 4C, Supplemental Data Set 3). However, the expression of senescence-associated TFs displayed similar dynamics in the SwAsp genotypes and therefore failed to explain the variation observed in their senescence timing (Fig. 4D). Evidently, like the major transcriptional patterns, the expression of individual senescence-associated TFs did not correlate with the onset of senescence across study years and genotypes. Instead, other factors – other gene expression patterns, hormone levels and genetic differences – could contribute more for the regulation of senescence onset.

**Figure 4.**
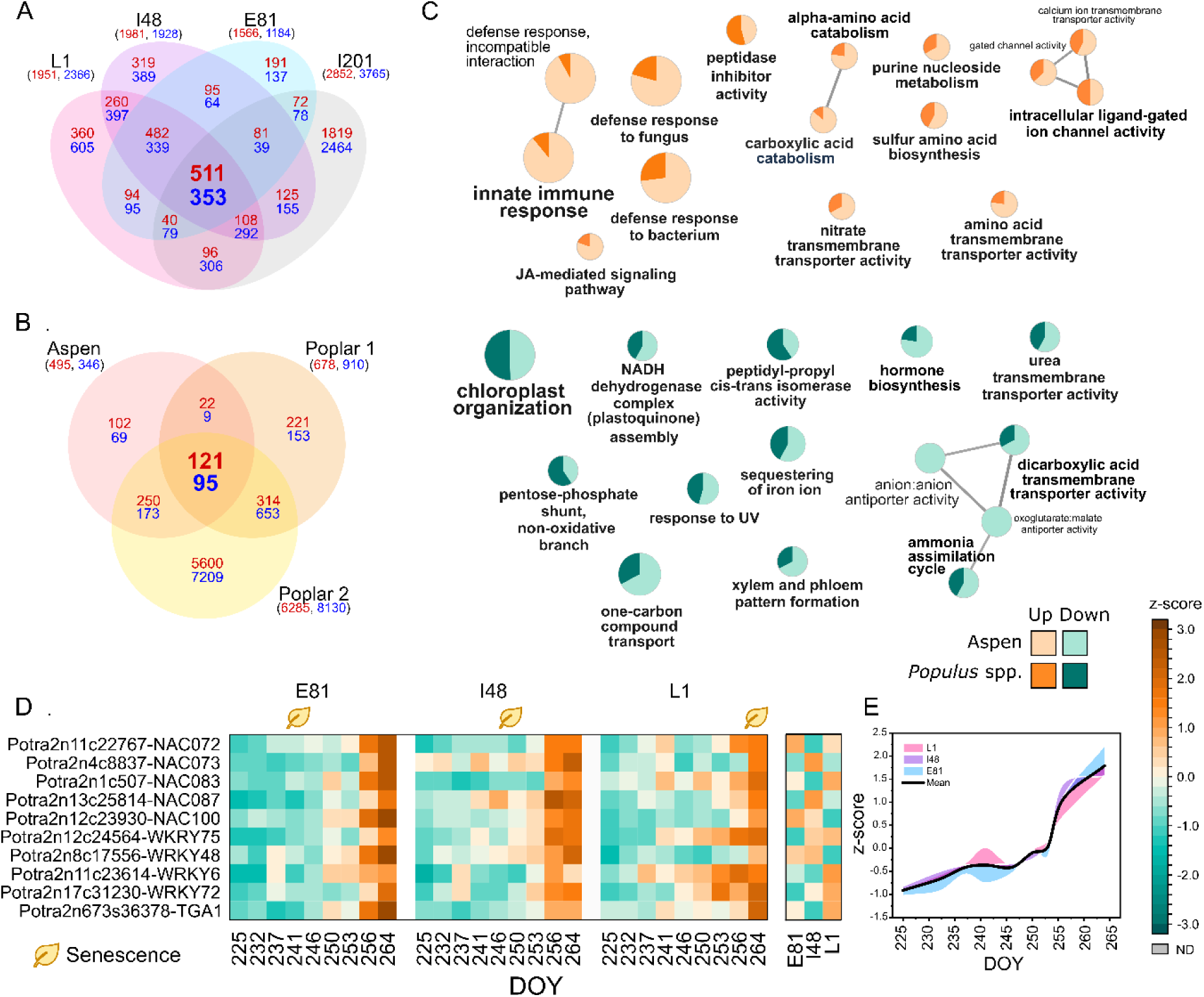
Senescence-associated genes (SAGs) in aspen (*Populus tremula*) and poplar (*P. trichocarpa)*. Venn diagram displays the overlap of up- and down-regulated SAGs in autumn in aspen (*P. tremula*, A) and in comparison with two data sets in poplar (*P. trichocarpa*) (B). For comparison, aspen gene IDs (Potra ID) were expressed as the best DIAMOND hit for *P. trichocarpa* gene IDs (Potri ID) obtained from PlantGenIE (https://plantgenie.org/). The lists of up- and down-regulated genes in poplar leaves were obtained from Leaf Senescence Database (LSD version 3.0, Li et al., 2020, Poplar 1) and from Lu et al., 2020 (Poplar 2). The number of up-regulated genes is in red and down-regulated in blue, shared genes between the data sets are in bold. The enriched Gene Ontology (GO) terms for biological processes of consistently up-regulated and down-regulated SAGs during autumn in aspen and in *Populus* spp. (C). The GO term nodes are coloured based on the proportion of shared genes in aspen and *Populus* spp. The expression patterns of senescence associated WRKY, NAC and TGA transcription factors displaying enhanced expression during autumn in three SwAsp genotypes (D, E). The data in the heatmap are mean expression normalized to z-score and the line graph depicts the expression pattern of the heatmap genes during autumn in each genotype relative to the mean, the deviation of genotype mean from the overall mean represented as a coloured area (mean of n=2-3 in each time point per genotype). Shared up- and down-regulated genes in autumn in aspen and in *Populus* spp. are in Supplemental Data Set 3.

### Autumn is accompanied by frequent stress events that can induce pro-senescence factors

Above and beyond the gradually changing transcriptional cascade (cascade-1), we detected another cascade (cascade-2) involving modules that displayed pronounced and often transient changes in their expression patterns during autumn (Fig. 3AB). In autumn 2018, modules in cascade-2 showed pronounced changes on and after 237 and 253 DOY correlating with short-term environmental fluctuations – seemingly coinciding with cloudy conditions that persisted over a few days (Fig. 3BC). Eigengene modules that showed either transient or sustained shifts in their expression in response to those weather shifts correlated with daily temperature variation, solar radiation, relative air humidity (RH%), and VPD (vapour pressure deficit) that all change concomitantly (Fig. 3BC, Fig. S9). In addition to the reduced amount of solar radiation and elevated air humidity (high RH and low VPD), the overcast days are accompanied by a shift in light quality with increased blue-to-red (B:R) and red-to-far red (R:fR) ratios of the light spectra; this was also true in the common garden (Fig. S16). This change in weather conditions affected a vast number of genes that affect multiple regulatory levels – transcriptional (RAV TFs, light signalling, circadian rhythm), post-transcriptional (RNA metabolism), epigenetic (histones, chromosome organisation) and translational (ribosome, translational initiation) levels.

Three modules associated with ribosome assembly and translation were induced in response to those weather shifts (module VI in cascade-1, VII in cascade-2) and one of them (module XX) showed more intensive up-regulation in early autumn in E81 and I48 than in L1 (Fig. 3B, Fig. S6). In line with the pronounced induction of genes encoding ribosomal proteins and translation initiation factors, endoplasmic reticulum (ER) stress and unfolded protein response (UPR)-related gene module was upregulated in mid-autumn in three SwAsp genotypes (module XVIII, Fig. 3B, Supplemental Data Set 1). Similarly, ER stress-related genes were upregulated in autumn in I201 in 2011 (modules 6, 13 and 16, Fig. 3A, Fig. S13).

The studied SwAsp genotypes varied considerably in their transcriptome responses between consecutive time points during autumn (Fig. 2B). Genes that displayed different response intensities, dynamics and relationships among the genotypes were the most interesting ones, since they could account for the differences in senescence onset. The more intensive shift in the transcriptome (PC7, Fig. 2B) and in the modules in cascade-2 in E81 and I48 than in L1 on 237 DOY (module XX, Fig. 3B) also reflected the enhanced expression of genes in downstream modules such as RAV TF regulated genes and genes involved in ethylene and abiotic stress responses, and programmed cell death (PCD, Fig. 5). Those included MAP kinases, WRKY and AP2/EREBP TFs such as *DREB1A/CBF3, RAP2*-*3* along with other stress-responsive genes such as *ZAT10 (ZINC FINGER PROTEIN 10), RDUF1* as well as a positive PCD regulator *LOL1* (*LSD1-LIKE 1*) (Fig. 5ABC, Supplemental Data Set 4). Other positive PCD regulators were also induced, whereas a negative PCD regulator *LSD1* (*LESION SIMULATING DISEASE 1*) was repressed, and these responses preceded senescence onset occurring earlier in genotypes with early-and intermediate-senescence and later in the season in the late-senescing genotype (Fig. 5ABC). Overall, the expression of positive PCD regulators increased during autumn and seemed to be amplified by the stress events (Fig. 5B). Their expression was also the highest in E81 and the lowest in L1 indicating that the genotypes had different expression of pro-death factors (Fig. 5AB).

**Figure 5.**
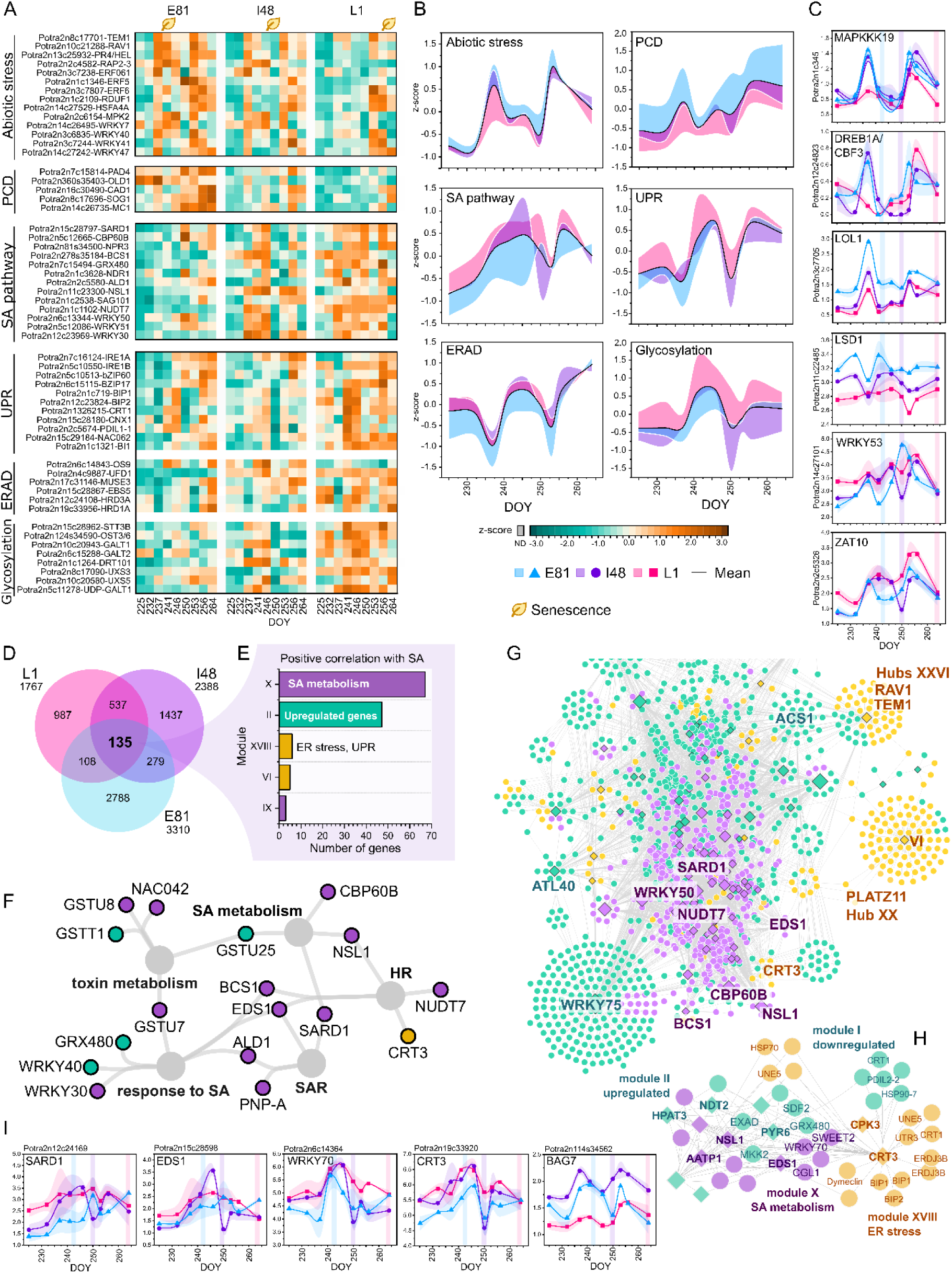
Genotypes show different dynamics of genes involved in ethylene and abiotic stress responses, programmed cell death (PCD),salicylic acid (SA)-mediated signalling pathway and endoplasmic reticulum (ER) stress responses. The expression patterns of genes involved in ethylene, abiotic stress, PCD, SA pathway and ER stress related processes in three SwAsp genotypes in autumn 2018 (A). Heatmap data are mean expression (n=2-3 in each time point per genotype) normalized to z-scores (A). Overall expression of the genes involved in the biological processes in each genotype expressed relative to the mean, deviation represented as a coloured area (B). The data of individual gene expression levels in the line graphs are VST- (variance stabilised transformation) normalized counts, mean ± SE (shadowed area), n=2-3 in each time point per genotype (C, I). The overlap of genes with positive correlation with SA levels in SwAsp genotypes (D). The assigned modules (E) and the Gene Ontology (GO) term network (F) of genes with positive correlation with SA levels in three SwAsp genotypes. Gene co-expression network displays the close neighbours of consistent SA-responsive genes (G) and a subnetwork (H) displays the potential mediators between temperature- and senescence-associated modules (modules I and II in cascade-1) and SA- (X) and ER stress-associated (XVIII) gene modules. Expression patterns of individual genes acting as potential regulators for and mediators between SA and ER stress responses (I). The network contains edges with correlation coefficient r > 0.6 and FDR adjusted *P*-value <0.05 common in the three genotypic networks. Nodes are coloured based on the associated cascade and the diamond shape displays genes with consistent positive correlation with SA levels. The lists of differently regulated genes in early autumn between genotypes are in Supplementary Data Set 4, extended GO term network of SA-responsive genes is in Fig. S17 and the details of the SA correlation results are in Supplementary Data Set 5. ERAD= ER associated protein degradation, UPR=unfolded protein response, SAR=systemic acquired resistance, HR=hypersensitive response

Similarly, abiotic stress-responsive genes such as *DREB/CBFs* and a module associated with cell death regulation were up-regulated before and at senescence onset in I201 (between 245-255 DOY, modules 11 and 16, Fig. 3A, Fig. S14). These results suggested that autumn was accompanied by weather shifts that caused stress for the trees especially in mid-autumn (237-253 DOY, hereafter defined as the *stress period*). This led to the re-wiring of the transcriptional network inducing ethylene, abiotic stress signalling and PCD genes (hereafter defined as the *pro-senescence phase*) at different times during autumn in the studied genotypes. Thus, the timing of this *pro-senescence phase* transition was dependent on the genotype and contributing to their senescence phenotype.

### Repression of salicylic acid-mediated transcriptional program coincided with senescence onset

In addition, both of the aforementioned cascades (cascade-1 and -2) were in the end interlinked with two large gene modules that displayed a transient repression coinciding with the start of rapid chlorophyll depletion (*senescence phase*) in the SwAsp genotypes (modules IX and X, Fig. 3B). The modules were enriched with genes involved in the regulation of salicylic acid (SA) metabolism and SA signalling pathway, leaf senescence, immune response, cell death, alternative respiration, macromolecule modification, nucleotide sugar metabolism and glycosylation (Fig. 3B, Supplemental Data Set 1). They also included several genes encoding receptors localised in the cell periphery at the interface of signal perception (Supplemental Data Set 1). Genes encoding membrane-bound proteins localised in the cell periphery and within the ER showed also different dynamics (significant time × genotype interaction) among the three SwAsp genotypes (Fig. 2D, Supplemental Data Set 4). The hub genes in the modules encoded positive regulators of the SA-mediated signalling pathway, systemic acquired resistance (SAR) and hypersensitive response (HR) such as two SARD1 (SAR DEFICIENT 1)-like TFs, calmodulin binding protein CBP60B (CALMODULIN-BINDING PROTEIN 60B) and EDS1 (ENHANCED DISEASE SUSCEPTIBILITY1) (Supplemental Data Set 1). The modules contained a large number of NAC and WRKY TFs such as *WRKY50, WRKY53* and *WRKY70* involved in SA/JA crosstalk (Fig. S11, Supplemental Data Set 1). Moreover, in the I201 network, *SARD1* acted as an important hub gene, up-regulated during autumn but transiently repressed at the onset of chlorophyll depletion (module 4, Fig. S14, Supplemental Data Set 2). These results suggested that in addition to the different timing of *pro-senescence phase* transition, the repression of the SA-mediated transcriptional program downstream of the two major cascades could play a key role in coordinating the start of chlorophyll depletion.

### Crosstalk between phytohormonal signalling pathways during the stress period can define senescence onset

Next, we profiled the levels of auxins (IAA), cytokinins (CKs), SA, ABA, and JA and their metabolites in the leaves of five SwAsp genotypes and investigated WGCNA eigengene module and gene expression relationship with phytohormone levels in three of them. Firstly, the different genotypes often had vastly different levels of several hormones and secondly, due to the strong short-time fluctuations, no single hormone level could be used as a sole reliable predictor for senescence phenotype. Not even IAA, that showed in general the lowest levels in early-senescing genotypes and the highest in the late-senescing ones (Fig. 6C). As a general pattern, the levels of active free forms of CKs and IAA decreased during autumn in all SwAsp genotypes, whereas the levels of their conjugated forms or catabolites increased (Fig. 6ABC). These metabolite levels correlated with the expression patterns of up- and down-regulated modules during autumn in the three SwAsp genotypes (cascade-1) indicating that along with the gradually decreasing air temperature the cascade was responding to CK and IAA metabolite levels (Fig. 3B, Fig. S9, S10 and S11). In line with this, the down-regulated module contained many auxin response factors (ARF) and AUX-IAA TFs (module I, Fig. S11C, Supplemental Data Set 1).

**Figure 6.**
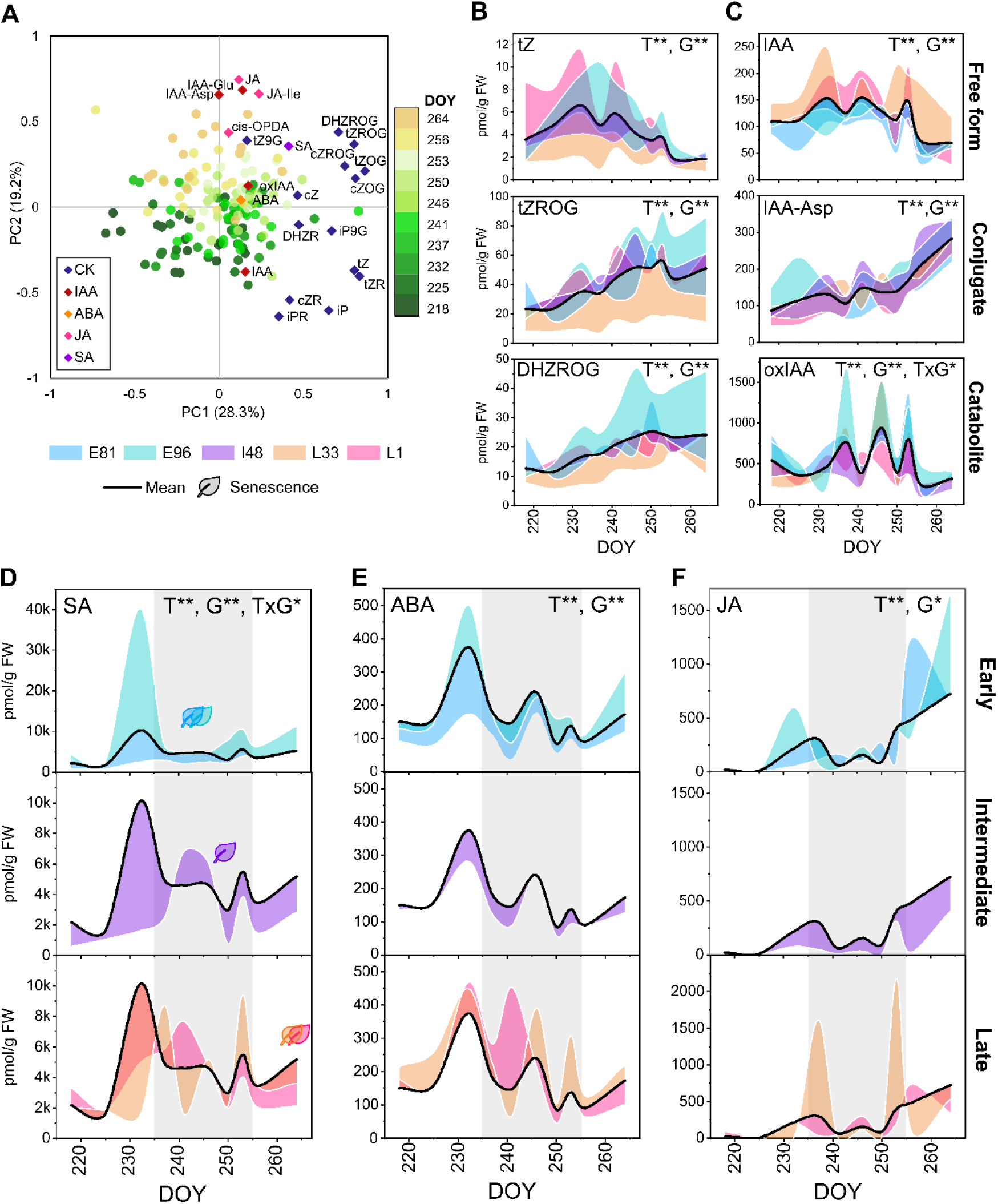
Phytohormone levels show pronounced temporal variation in autumn reflecting different responses to stress events between aspen genotypes. Overall variation in the phytohormone profile in autumn 2018 is depicted in the bi-plot of principal component analysis and samples are coloured based on the time points (PCA, A). The levels of trans-zeatin (cytokinin, CK) metabolites (B), auxin (IAA) metabolites (C), salicylic acid (D), abscisic acid (E) and jasmonic acid (F) in the leaves of five SwAsp genotypes in autumn 2018. The grey area represent the stress period during autumn. The effect of time (T, nine time points 218-264 DOY) and genotype (G) and their interaction (T×G) were tested with two-way ANOVA (FDR adjusted P-value <0.01**, <0.05*). Metabolite levels in SwAsp leaves are expressed relative to the mean, deviation represented by a coloured area (mean of n=2-3 in each time point per genotype). See details of statistical results and the levels of all phytohormones in autumn 2018 in Supplementary Data Set 5 and Fig. S8, respectively.

The levels of IAA catabolite (oxIAA) and ABA, JA and SA levels varied markedly during autumn. ABA and oxIAA levels were transiently elevated on three days in all genotypes reflecting frequent stress events (Fig. 6CDEF). JA levels were elevated during autumn and SA levels were upregulated at different times in autumn among the genotypes (significant genotype × time interaction effect, Fig. 6D). On several occasions elevated SA levels were accompanied by elevated levels of other stress-induced phytohormones, JA and/or ABA, that are antagonistic to SA signalling (Fig. 6DEF). ABA levels during autumn did not correlate with changes in the leaf transcriptome (Fig. S10). Overall, very few genes known to be involved in the metabolism of stress-induced hormones (ABA, JA) in Arabidopsis displayed a consistent relationship with the associated metabolite levels in aspen leaves, except for SA (Supplemental Data Set 5).

We confirmed that SA levels correlated positively with the expression of *SARD1* and *CBP60B* (hub genes in SA-associated module X) involved in SA biosynthesis, and other genes involved in SA-mediated SAR and HR, innate immunity, and leaf senescence, evidently defining the transcriptional response in the three SwAsp genotypes (Fig. 5F, Supplemental Data Set 5). In addition, SA levels correlated positively with the expression of *NCED3* encoding 9-cis-epoxycarotenoid dioxygenase, a key enzyme in the ABA biosynthesis, with *ACS1* encoding 1-aminocyclopropane-1-carboxylate synthase involved in ethylene biosynthesis, and with genes involved in SA/JA crosstalk suggesting a complex relationship among these hormones (Fig. 5G, Supplemental Data Set 5). In general, SA levels during the *stress period* were lower in early-senescing genotypes than in the other genotypes that senesced later (Fig. 6D). Therefore, in line with the transcriptional program associated with SA metabolism and SA/JA-crosstalk, whose temporal pattern matched with the timing of senescence onset, SA appeared as the best candidate for defining the timing of senescence.

### Alleviated proteotoxic stress and redox homeostasis in autumn can affect senescence onset

Finally, as the SA-mediated signalling pathway potentially could modulate senescence timing, a gene co-expression network was constructed to investigate the connection of SA-responsive genes within the transcriptional network. To focus on consistent relationships, the overlap of the three genotypic co-expression networks was visualised (Fig. 5GH). Interestingly, SA levels and SA-associated gene modules showed a connection with the ER stress-associated genes and gene modules with a stronger positive relationship in L1 and I48 that displayed a pronounced increase in SA levels during the *stress period* in early autumn, and a weaker relationship in E81 that did not (Fig. S10, Fig. S17). Accordingly, SA levels correlated with the expression of genes involved in protein modification, glycoprotein and nucleotide sugar metabolism and with genes encoding ATL family RING-H2 ubiquitin ligases linked to defence responses and ER-associated degradation of misfolded proteins by ubiquitination (ERAD pathway) (Fig. 5GH, Supplemental Data Set 5).

Moreover, co-expression network analysis identified genes accounting for glycoprotein quality control in ER (ERQC) by the chaperone system as consistent mediators controlling the information flow through the network structure – between SA- and ER-stress-associated gene modules. Calreticulin 3 (*CRT3*) that was assigned into the ER stress and UPR module (module XVIII) correlated positively with SA levels and was co-expressed with SA-pathway genes: *EDS1, WRKY70* and *NSL1 (NECROTIC SPOTTED LESIONS 1* (Fig. 5HI). On the other hand, genes connecting the ER stress module with the cascade-1 (module II) included *SDF2* (*stromal-cell derived factor 2-like*) that forms an ERQC complex with ERDJ3B and BIP1 (Fig. 5H). Together these correlation and co-expression analyses displayed a complex regulatory network during autumn where SA-mediated (module IX and X) and temperature-responsive and senescence-associated programs (cascade-1: module I and II) were interlinked to modulate ER related functions mainly via nucleotide sugar and glycoprotein metabolism as well as the quality control system of glycoproteins (Fig. 5H), many of which are membrane-bound receptors. Furthermore, the expression of several SA-, UPR-, ERAD- and glycosylation-related genes was the lowest in E81 and strongly repressed in I48 in early and mid-autumn potentially accounting for their senescence onset due to hindered stress tolerance (Fig. 5AB, Supplemental Data Set 4). *BAG7* was an exception as it showed higher expression in E81 and I48 than in L1 in early autumn (Fig. 5I). Furthermore, the genes encoding Plant Natriuretic Peptide A (PNP-A), alternative oxidases (AOXs), nudix hydrolases (NUDT2, NUDT7) and glutathione transferases (GSTs) were co-expressed with SA-pathway genes and/or correlated positively with SA levels among other genes involved in ROS metabolism and cellular respiration (Fig. 5FG, Fig. S17, Supplemental Data Set 5) suggesting that the activated SA signalling pathway promoted also cellular redox homeostasis and the activation of retrograde signalling cascades potentially affecting energy metabolism.

Our findings were further supported by the co-expression network analysis in genotype I201 in autumn 2011. A module at the end of the hierarchical transcriptional network contained genes involved in metabolic pathways and encoding receptor-like proteins (hubs). The modules directly upstream were enriched with genes involved in ER stress responses, glycosylation and cell death regulation that were connected to the SA-associated gene module and the two transcriptional cascades (Fig. 3A, Fig. S13, Supplemental Data Set 2). Thus, in accordance with SA levels, the SA-mediated transcriptional program appeared as the best candidate for coordinating the timing of leaf senescence in aspen with its connection to the two cascades, cell death regulation, other stress-induced phytohormones and to the post-translational processes that contribute to metabolic homeostasis, proteome maintenance and proteotoxic stress tolerance. Considering all the presented data, we propose that first the decision to senesce is defined by the trees’ response to environmental factors (cascade-1 and - 2), and that the start of rapid chlorophyll depletion is fine-tuned by SA signalling pathway that can counteract the onset by alleviating the evoked metabolic stress.

## DISCUSSION

There must be a trigger for the spectacular phenomenon of autumn senescence but what is it? These data, together with our published results (Jansson and Thomas, 2008, Edlund et al., 2017) have led us to a suggestion that the way plant scientists traditionally look at autumn leaf senescence could be misleading. There is a remarkable complexity in how aspen leaves enter senescence, including within the same tree individual (Lihavainen et al., 2020, Fataftah et al., 2021). It is known that many internal and external factors can induce senescence, and that abiotic stress, immunity responses, cell death and senescence share many components in their signalling network (Bruggeman et al., 2015). Immunity and cell death responses are affected by temperature, light and humidity conditions (Yoshioka et al., 2001, Huang et al., 2010, Chai et al., 2015), that all change in parallel in autumn. Any of the challenging environmental factors in autumn such as cold or change in light conditions could induce signals that will open “the gateway to senescence” creating a situation where *“all roads lead to Rome”*. This hypothesis could explain why we found patterns in our analyses that indicated that trees may not rely on only one pathway to properly time their autumn senescence. We found that autumn was accompanied by multiple stress events that could in theory cause the trees to senescence and that two tightly interlinked transcriptional cascades appeared to respond to environmental variation and affect senescence onset via the SA-mediated pathway and downstream metabolic stress responses that can be evoked by various factors.

Although we could not identify any single gene or hormone marker that would change drastically and explain well why some aspens enter senescence early and others late, we identified patterns and gene modules closely associated with the changes in weather conditions, and a sequence of changes at hormone and transcriptional levels and their connection with post-transcriptional processes associated with the regulation of the onset and progression of senescence in response to autumnal cues under natural conditions (Table S1). Moreover, many of the biological processes and associated transcriptional programs that have been previously linked to the regulation of senescence and cell death in controlled laboratory studies with plant and animal models (Woo et al., 2013, Woo et al., 2019) change in parallel and are tightly coordinated during autumn in aspen in a complex manner. The senescence transcriptome has been studied in, for example, Arabidopsis (Balazadeh et al., 2008; Breeze et al., 2011), tobacco (Li et al., 2017), poplar (*P. trichocarpa*, Lu et al., 2020, *P. tomentosa* Wang et al., 2021) and red maple (*Acer rubrum*, Zhu et al., 2020), and these studies have led to the identification of thousands of SAGs that are induced or repressed during autumn and during developmental or stress-induced senescence (Leaf senescence database, LSD 3.0, Li et al., 2020). Our meta-analysis of transcriptome alterations during autumn in aspen and poplar (Li et al., 2020, Lu et al., 2020) showed that the most conserved responses in *Populus* leaves during autumn were the repression of chloroplast processes, and the induction of immune and defence responses mediated by NAC and WRKY TFs; TF families were well represented among SAGs. However, the induction or repression of SAGs alone could not explain the onset of autumn senescence in aspen under natural conditions.

Our results also suggest that there may be a link between chromatin remodelling and gene expression patterns in autumn. Indeed, epigenetic mechanisms such as histone modification and DNA methylation have been associated with dormancy regulation in trees (Karlberg et al., 2010, Cooke et al., 2012), and with the regulation of plant immunity and the SA pathway (Ramirez-Prado et al., 2018, Chen et al., 2020) and leaf senescence through SAG expression in Arabidopsis (Ay et al., 2014; Wang et al., 2019). The potential role of epigenetic mechanisms in the regulation of autumn senescence, for example by affecting the ability of trees to respond to external and internal cues, remains an intriguing topic for further study.

It is noteworthy that the gene modules containing SAGs and chlorophyll metabolism genes correlated with chlorophyll levels if senescence onset occurred in early or mid-autumn, but not if senescence was initiated late in the season. Perhaps SAGs and chlorophyll metabolism genes (in cascade-1) contribute to adjusting the rate of senescence, after it has been initiated, to ensure efficient nutrient resorption before frost kills the leaf cells, in line with the observation that cold temperature accelerates the rate of leaf senescence once it has been initiated (Fracheboud et al., 2009). Milder temperatures could give a slower senescence progression in early autumn, and more rapid progression in late autumn when the expression of SAGs and the levels of positive (JA) and negative senescence regulators (CKs and IAA), have already changed; in other words, senescence is an active process and the leaves also respond to environmental factors after it has been induced.

Many of our observations connect senescence with abiotic stress responses, and an overlap in the regulation of plant immunity responses and the senescence network has been shown previously (Guo and Gan, 2012, Zhang et al., 2020). Stress-induced phytohormones and stress-responsive genes are typically regarded to positively regulate senescence and are considered as SAGs since their levels or expression are elevated during stress-induced and developmental senescence in other plant species (Buchanan-Wollaston et al., 2005; Balazadeh et al., 2008; Breeze et al., 2011; Guo and Gan 2012). Here we show that the onset of autumn senescence in aspen was not defined by high expression of SAGs or high levels of ABA or JA. Our data suggest that senescence may be conditional on the activation of ethylene/abiotic stress and/or PCD signalling pathways after which the timing and progress of senescence is fine-tuned by the interplay of temperature- and senescence-associated (cascade-1) and SA-mediated transcriptional programs where SAGs are integral nodes.

We note that in the *pro-senescence phase*, genes encoding RAV TFs that are involved in the regulation of ethylene signalling, senescence and the photoperiodic pathway (Woo et al., 2010; Matías-Hernández et al., 2014) were induced. Furthermore, *LOL1* (*LSD1 like 1*), involved in the initiation of ROS-mediated PCD, was induced, whereas antagonistic *LSD1* (Epple et al., 2003; Valandro et al., 2020) was slightly repressed, indicating a shift in the expression of pro-death relative to pro-survival factors. Moreover, the expression of pro-death factors was initially different between the genotypes (high in the early-senescing ones), potentially accounting for their senescence phenotypes. Indeed, LSD1 regulates cell death in response to temperature and light cues (Rustérucci et al., 2001, Huang et al., 2010, Wituszyńska et al., 2015) and the runaway cell death phenotype in *lsd1* mutant is SA- and EDS1-dependent (Rustérucci et al., 2001, Aviv et al., 2002) with a potential to integrate the PCD regulon with environmental signals and with the SA-mediated signalling pathway.

After the *pro-senescence phase*, the shift into the *senescence phase* seemed to coincide with the de-regulation of the SA-mediated transcriptional program driven mainly by SARD1 that in other species promotes SA biosynthesis and regulates signalling downstream of immune receptors to induce SAR or HR (Zhang et al., 2010; Sun et al., 2015; Zhou et al., 2018). Overall, in aspen leaves, SA seems to delay senescence, not promote it as is often observed in Arabidopsis (Morris et al., 2000; Vogelmann et al., 2012). A dual role of SA in regulating senescence and PCD has been established in several recent reports (*e*.*g*., Radojičić et al., 2018; Lee et al., 2020) and SA can delay senescence by activating autophagy (Yin et al., 2020, Rigault et al., 2021) or by alleviating ER stress by modulating glycosylation of proteins and metabolites (Huang et al., 2018; Lai et al., 2018; Lee et al., 2020). For example, BiP proteins (ER chaperones) can act as positive or negative cell death regulators depending on the activation of SA-mediated HR (Carvalho et al., 2014). Moreover, SA pathway has a dual role in regulating cellular ROS production by either supressing it (Herrera-Vásquez et al., 2015, Poór 2020), or by amplifying ROS signals such as those observed during senescence (Guo et al., 2017). Overall, our network analyses support the understanding that the activated SA signalling pathway promotes cellular redox homeostasis in autumn.

Our data also suggest a connection between the SA pathway and unfolded protein response (UPR) that alleviates ER stress or induces cell death while prolonged (Schäfer and Eichmann, 2012, Afrin et al., 2020). The expression of UPR genes either correlated with SA levels or were co-expressed with SA pathway genes, especially in the genotypes that induced the SA pathway during autumn. It is unclear how SA signalling activates UPR in response to ER stress (Nagashima et al., 2014; Poór et al., 2019), but our data suggest that *WRKY70, EDS1, NSL1* and *CRT3* along with genes involved in protein degradation (ERAD), nucleotide sugar metabolism and glycoprotein quality control may connect the SA pathway and ER stress regulon in aspen leaves. Protein glycosylation requires nucleotide sugars, and the OST complex is the main enzyme driving N-glycosylation in the ER (Deng 2013; Howell 2013). Calnexins (CNXs) and calreticulins (CRTs) are molecular chaperones that participate in protein folding and quality control of glycoproteins (Pattison and Amtmann, 2009; Howell 2013), many of which are receptor proteins translocated via trans-Golgi network to the plasma membrane. In Arabidopsis, WRKY70 negatively regulates developmental senescence and acts at the convergence point of antagonistic phytohormone pathways (inducing SA and repressing JA) (Li et al., 2004, Ülker et al., 2007), EDS1 is involved in ROS- and SA-dependent cell death (Rustérucci et al., 2001), and CRT3 is linked to the maturation of receptor-like kinases (RLKs) that affect cell death (Vitale et al., 2009, Sun et al., 2014). Interestingly, *BAG7* expression was enhanced in early autumn in E81 and I48 and BAG (Bcl-2-associated athanogene) family proteins have been implicated to regulate leaf senescence and proposed to function as potential “switches” between cell death and cell survival (Li et al., 2016; Ren et al., 2021).

In Figure 7, we have, based on the arguments above, made a tentative model to explain the senescence phenotypes: fast senescence onset after the *pro-senescence phase* in early autumn (E81, E96), delayed senescence onset after the *pro-senescence phase* in mid-autumn (I48, I201) or no senescence onset until the *pro-senescence phase* is reached in late autumn (L1, L33). We suggest that the timing of transcriptional and hormonal responses is crucial for the regulation of senescence onset. The early-senescing genotype E81 showed the highest expression of positive PCD regulators, and only minor or no induction of SA pathway genes or SA levels in mid-autumn during the *stress period* after the *pro-senescence phase* potentially contributing to the early onset. In the intermediate-senescing genotypes (I48, I201), the SA pathway was activated after the *pro-senescence phase* and it remained active during the *stress period* for ca. one week until it was repressed, and senescence was initiated. If the changes in environmental conditions in early autumn did not induce ethylene-, abiotic stress- and PCD-related genes, the following *stress period* did not trigger senescence onset. Instead, the genotype displayed high SA levels (and other stress-induced hormones), high expression of SA pathway-, glycosylation-, UPR- and ERAD-related genes that could provide stress tolerance by restoring ER and redox homeostasis. Only after the expression of PCD, ethylene and abiotic stress genes was elevated in late autumn did the SA levels and the expression of SA pathway genes decrease, and senescence was initiated. By then SAG expression had changed and the levels of positive (JA) and negative hormonal regulators (CK, IAA) were elevated and depleted, respectively, promoting senescence. After growth cessation and bud set, senescence onset may be defined by at least two layers of regulation in a conditional manner; the up-regulation of pro-senescence factors in response to environmental variation may prime the trees for senescence, and the de-regulation of SA-mediated stress tolerance mechanisms destabilise ER function and cellular redox status and potentiate the pro-senescence signals, resulting in senescence onset and developmental PCD.

**Figure 7.**
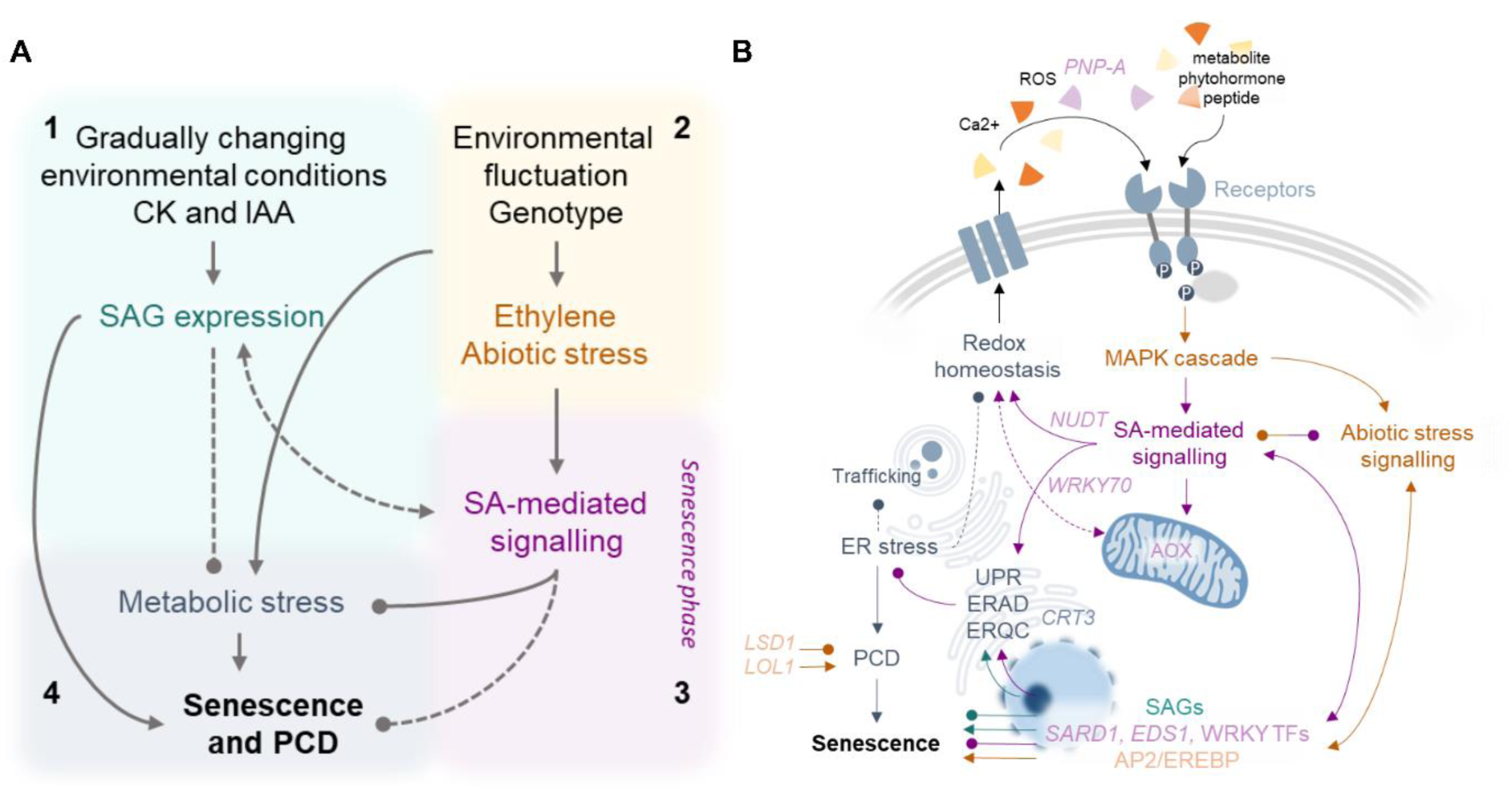
Autumn senescence is coordinated by two parallel transcriptional regulatory cascades in response to environmental signals and phytohormones. A hypothetical model displays the two main transcriptional cascades acting in parallel during autumn, one 1) influenced largely by gradually decreasing air temperature and the levels of cytokinin (CK) and auxin (IAA) metabolites, similar in aspen genotypes, and another one 2) that shows genotype-dependent rewiring of in response to short-term environmental fluctuation inducing ethylene, abiotic stress and PCD gene expression. Both cascades are connected to 3) the salicylic acid (SA)-mediated signalling pathway and 4) metabolic stress responses (such as ER stress, unfolded protein response-UPR) that appear to play key roles in fine-tuning the timing of senescence onset under challenging autumn conditions. A hypothetical model of the connection between biological processes involved in the regulation of the timing of autumn senescence (B). The different sensitivity to external and internal cues between the genotypes can relate to the different expression of genes (significant time × genotype interaction and induced/repressed in response to environment and phytohormones) encoding receptors at the interface of signal perception. Furthermore, ER stress during autumn can hinder the maturation and establishment of receptor proteins. The SA signalling pathway, that can antagonize ABA/JA/ET-mediated signalling, appeared to affect also cellular respiration and redox status by mediating the expression of *PNP*-*A*, alternative oxidase (*AOXs*) and nudix hydrolase (*NUDT*) genes. Temperature-responsive and senescence-associated transcriptional program (cascade-1), the ER stress and UPR regulon (in cascade-2), and the SA-mediated pathway were connected via protein glycosylation and protein folding via ER-quality control (ERQC) system. Hence, we propose a model where autumn senescence onset in aspen is conditional to the activation of pro-senescence factors in response to environmental/metabolic stress and the timing of chlorophyll depletion is fine-tuned by the SA-mediated transcriptional program that affects stress tolerance mechanisms (UPR, redox metabolism, phytohormone antagonisms) and thus senescence onset. The de-regulation of pro-survival mechanisms can disrupt cell functioning and potentiate pro-death factors to initiate autumn senescence and developmental PCD. ERAD=Endoplasmic reticulum associated degradation

Autumn is accompanied by ample weather shifts and rapid variation in the diurnal light regime leading to many responses in aspen leaves. We are convinced that these profound changes in the leaf transcriptome and hormone profiles that are evoked by these variations are important for senescence regulation, even though they may not directly explain the molecular mechanism initiating the process. The expression of genes encoding membrane-bound receptors in the cell periphery and in ER displayed significantly different dynamics between aspen genotypes and were induced and repressed in response to weather shifts, and hormone levels and potentially the receptor proteins failed to maturate due to ER stress. Perhaps the differences in the perception and transduction of signals via receptor proteins and signalling cascades can explain genotypic sensitivity to environmental cues and metabolic signals such as ROS, calcium and hormones and account for their senescence phenotype.

We would like to stress the importance of not only studying leaf senescence under controlled conditions but also under field conditions to which plants are adapted. Conclusions made by associations, correlations or even mutant studies that cannot be corroborated by studies on plants growing under natural conditions are likely to overestimate the importance of the conclusions. Finally, one may question if “omics profiling” can be useful for understanding senescence, despite that in our case it has not has made us understand how the process is initiated at molecular level. One obvious possibility is that the key internal changes which are triggered by external factors are downstream of mRNA levels and hormones, for example on a translational, or post-translational level, or that it is a purely metabolic signal. We are currently also investigating this with the very same setup, and this will be the subject of coming publications. Nevertheless, efforts like this also give ideas to follow up, in our case for example the involvement of the SA pathway, ROS, receptor proteins and proteotoxic stress in autumn senescence regulation. Integration of multi-omics techniques combined with screening of physiological traits has a great potential to generate novel hypotheses to decipher the complex regulation underlying senescence (Großkinsky et al., 2018) and may eventually enable the establishment of a diagnostic set of molecular markers for autumn senescence in trees.

To conclude, after comparing the mRNA and hormone levels and co-expression network structures between years and genotypes with different senescence timing under the same (fluctuating) environmental conditions, we could not identify a “master switch” *i*.*e*., a single gene or hormone marker that could be used as a sole predictor for the senescence phenotype. Instead, our results demonstrate that more than one transcriptional cascade contribute to senescence regulation. We suggest that first the re-wiring of the transcriptional regulatory network in response to environmental factors induces pro-death factors marking the transition into the *pro-senescence phase* that primes the trees for senescence initiation. Afterwards the timing of senescence onset seems to be fine-tuned by the SA signalling pathway that can alleviate metabolic stress. Overall, our results suggest that the phenotypic differences in senescence onset are likely linked to the different timing of cellular responses between aspen genotypes, and to their intrinsic capacity to activate pro-survival mechanisms under stressful autumn conditions. In other words, the onset of autumn senescence appears to be defined by the shift in balance between pro-death and pro-survival factors in response to environmental and internal metabolic cues.

## ACKNOWLEDGMENTS

This research has been supported by funding from the Swedish Research Council VR, Formas, Kempestiftelserna, Swedish Foundation for Strategic Research (SSF), Knut and Alice Wallenberg foundation, Vinnova,Trees for the Future (T4F) project and SE2B Horizon 2020 We thank the UPSC bioinformatics facility for their support in transcriptome data processing, Swedish Metabolomics Centre (SMC) for their support in phytohormone analysis and Skogforsk in Sävar for maintaining the SwAsp and UmAsp collections.

## AUTHOR CONTRIBUTIONS

JL, NF, KMR and SJ planned the experiments, JL, NF and KMR collected the samples and performed the measurements in the field, JL, PB and JŠ performed the experiments, JL and ND processed and analyzed the transcriptomics data, JL, JŠ and ON performed the phytohormone analysis, JL and PB analyzed the physiological data, JL and SJ wrote the paper and all authors contributed to the integration of the results and revision of the paper.

## DATA AVAILABILITY

Transcriptome data has been deposited at the European Nucleotide Archive (ENA, https://www.ebi.ac.uk/ena/browser/home) under the accession number: PRJEB51801. Transcriptome data for I201 in 2011 are available in Gene Expression Omnibus (GEO) repository with accession number GSE86960. Other data are available within the article and its supplementary material.

## SUPPLEMENTAL DATA

**Table S1**. The variation in the transcriptome data in autumn 2018 as explained by environmental factors, genotype and phytohormone levels.

**Figure S1**. Autumn senescence onset in three Swedish aspen (SwAsp) genotypes in autumn 2011 and clonal trees of I201 in autumn 2018 in the common garden.

**Figure S2**. Autumn senescence onset and rate in three Swedish aspen (SwAsp) genotypes in controlled greenhouse conditions.

**Figure S3**. Time-dependent patterns of significant principal components in principal component analysis (PCA) of global transcriptome data in three aspen genotypes in autumn 2018.

**Figure S4**. Score plot (A) and time-dependent patterns of principal components (B) in principal component analysis (PCA) of global transcriptome data in aspen genotype I201 in autumn 2011.

**Figure S5**. Weighed gene co-expression network analysis (WGCNA) statistics of 2018 data.

**Figure S6**. Preservation of WGCNA eigengene modules and module relationships in three SwAsp genotypes in 2018.

**Figure S7**. Weather parameters in autumn 2011 and 2018 in Umeå, Sweden.

**Figure S8**. The levels of phytohormones in the leaves of five SwAsp genotypes in autumn 2018.

**Figure S9**. Genotypic and consensus correlation plots of signature expression of WGCNA eigengene modules with weather parameters in autumn 2018.

**Figure S10**. Genotypic and consensus correlation plots of signature expression of WGCNA eigengene modules with metabolic markers in autumn 2018.

**Figure S11**. Correlation network displaying relationships between WGCNA eigengene modules, weather parameters and phytohormone levels in three SwAsp genotypes in autumn 2018 (A). Eigengene patterns (B), the abundance of transcription factor families and chromatin remodelling factors in each gene module (C) and the number of modules with significant correlation with environmental factors and phytohormone levels in consensus data and in each genotype (D).

**Figure S12**. Weighed gene co-expression network analysis (WGCNA) statistics of 2011 data.

**Figure S13**. Weighed gene co-expression network analysis (WGCNA) eigengene patterns and the expression of selected genes in I201 in autumn 2011.

**Figure S14**. Correlation plot of signature expression of WGCNA eigengene modules with weather parameters and chlorophyll content index (CCI) in I201 in autumn 2011.

**Figure S15**. Expression patterns of chlorophyll metabolic genes and their correlation with chlorophyll content index (CCI).

**Figure S16**. Solar spectral quality in autumn 2020 in Umeå, Sweden.

**Figure S17**. Gene Ontology (GO) term enrichment analysis of salicylic acid-responsive genes in three SwAsp genotypes in autumn 2018.

## Supplemental Methods

**Supplemental Data Set 1 Table a-k**. Weighed gene co-expression analysis (WGCNA) results in three SwAsp genotypes in autumn 2018. Module membership (a), correlation with environmental parameters (b) and phytohormone levels (c). The number of genes with significant correlation with weather parameters and metabolic markers in each genotype and consensus network (d). The number and relative abundance (%) of transcription factors (e), chromatin remodelling factors (f) and senescence-associated genes (SAGs, g) in each module. The top 20 hub genes in the modules (h). Enriched Gene Ontology (GO) terms (I, j, k) and Kyoto Encyclopedia of Genes and Genomes (KEGG) pathways (l) in each module. Eigengene scores of samples (m).

**Supplemental Data Set 2 Table a-i**. Weighed gene co-expression network analysis (WGCNA) results in genotype I201 (campus tree) in autumn 2011 (a). The number of genes with significant correlation with weather parameters and chlorophyll content index (b). The number and relative abundance (%) of transcription factors (c) and chromatin remodelling factors (d) in each module. The top 20 hub genes (e) and enriched Gene Ontology (GO) terms (f,g) and Kyoto Encyclopaedia of Genes and Genomes (KEGG) pathways in each module (h). Eigengene scores of samples (i).

**Supplemental Data Set 3 Table a-d**. Consistently up- or down-regulated genes (Senescence-associated genes-SAGs) during autumn in aspen (a) and in *Populus* spp. (b). The number of SAGs with significant correlation with weather parameters and metabolic markers in each genotype and in consensus data (c). Pfam (protein domains) of consistently up- or down-regulated genes in *Populus* spp. during autumn (d).

**Supplemental Data Set 4 Table a-e**. Differentially expressed (DE) genes as affected by time, genotype and time × genotype interaction in SwAsp genotypes in autumn 2018 (a). The list of genes with significant interaction effect (b) and enriched Gene Ontology (GO) terms and Pfam (protein domain) in the gene set (c). Comparison of DE genes between the SwAsp genotypes on different time points in early and mid-autumn (237-250 DOY, d-e).

**Supplemental Data Set 5 Table a-j**. The list of identified phytohormones (a) and their levels in the leaves of five SwAsp genotypes in autumn 2018 (b). The effects of time, genotype and their interaction on metabolite levels (c). The list of genes with significant correlation with leaf salicylic acid (SA) levels in autumn 2018 in individual SwAsp genotypes and in the consensus data (d-e). The number of SA-responsive genes in the gene modules (f) and their correlation with environmental parameters in aspen genotypes (g). The list of genes displaying consistent correlation with abscisic acid (ABA, h) and jasmonic acid (JA, i) levels in three SwAsp genotypes in 2018. The expression patterns of genes involved in phytohormone metabolism (based on Arabidopsis) in aspen genotypes in autumn 2011 and 2018 (j).

**Supplemental Data Set 6**. Correlation between the expression of genes involved in chlorophyll metabolism, weather parameters and chlorophyll content index (CCI) in aspen genotypes in autumn 2011 and 2018.

## REFERENCES

Afrin, T., Diwan, D., Sahawneh, K., & Pajerowska-Mukhtar, K. (2020). Multilevel regulation of endoplasmic reticulum stress responses in plants: where old roads and new paths meet. Journal of experimental botany, 71(5), 1659–1667.

Aviv, D. H., Rustérucci, C., Iii, B. F. H., Dietrich, R. A., Parker, J. E., & Dangl, J. L. (2002). Runaway cell death, but not basal disease resistance, in lsd1 is SA-and NIM1/NPR1-dependent. The Plant Journal, 29(3), 381–391.

Ay, N., Janack, B., & Humbeck, K. (2014). Epigenetic control of plant senescence and linked processes. Journal of experimental botany, 65(14), 3875–3887.

Balazadeh, S., Riaño-Pachón, D. M., & Mueller-Roeber, B. (2008). Transcription factors regulating leaf senescence in Arabidopsis thaliana. Plant biology, 10, 63–75.

Bindea, G., Mlecnik, B., Hackl, H., Charoentong, P., Tosolini, M., Kirilovsky, A., … & Galon, J. (2009). ClueGO: a Cytoscape plug-in to decipher functionally grouped gene ontology and pathway annotation networks. Bioinformatics, 25(8), 1091–1093.

Bindea, G., Galon, J., & Mlecnik, B. (2013). CluePedia Cytoscape plugin: pathway insights using integrated experimental and in silico data. Bioinformatics, 29(5), 661–663.

Bolger, A. M., Lohse, M., & Usadel, B. (2014). Trimmomatic: a flexible trimmer for Illumina sequence data. Bioinformatics, 30(15), 2114–2120.

Breeze, E., Harrison, E., McHattie, S., Hughes, L., Hickman, R., Hill, C., … & Buchanan-Wollaston, V. (2011). High-resolution temporal profiling of transcripts during Arabidopsis leaf senescence reveals a distinct chronology of processes and regulation. The Plant Cell, 23(3), 873–894.

Bruggeman, Q., Raynaud, C., Benhamed, M., & Delarue, M. (2015). To die or not to die? Lessons from lesion mimic mutants. Frontiers in plant science, 6, 24.

Buchanan-Wollaston V, Earl S, Harrison E, Mathas E, Navabpour S, Page T, Pink D (2003) The molecular analysis of leaf senescence—a genomics approach. Plant Biotechnol J 1:3–22

Buchanan-Wollaston V, Page T, Harrison E, Breeze E, Lim PO, Nam HG, Lin JF, Wu SH, Swidzinski J, Ishizaki K, Leaver CJ (2005) Comparative transcriptome analysis reveals significant differences in gene expression and signalling pathways between developmental and dark/starvation-induced senescence in Arabidopsis. Plant J 42:567–585

Carvalho, H. H., Silva, P. A., Mendes, G. C., Brustolini, O. J., Pimenta, M. R., Gouveia, B. C., … & Fontes, E. P. (2014). The endoplasmic reticulum binding protein BiP displays dual function in modulating cell death events. Plant physiology, 164(2), 654–670.

Chai, T., Zhou, J., Liu, J., & Xing, D. (2015). LSD1 and HY5 antagonistically regulate red light induced-programmed cell death in Arabidopsis. Frontiers in Plant Science, 6, 292.

Chen, J., Clinton, M., Qi, G., Wang, D., Liu, F., & Fu, Z. Q. (2020). Reprogramming and remodeling: transcriptional and epigenetic regulation of salicylic acid-mediated plant defense. Journal of experimental botany, 71(17), 5256–5268.

Cooke, J. E., Eriksson, M. E., & Junttila, O. (2012). The dynamic nature of bud dormancy in trees: environmental control and molecular mechanisms. Plant, cell & environment, 35(10), 1707–1728.

Delhomme, N., Mähler, N., Schiffthaler, B., Sundell, D., Mannapperuma, C., Hvidsten, T. R., & Street, N. R. (2014). Guidelines for RNA-Seq data analysis. Epigenesys protocol, 67, 1–24.

Deng, Y., Srivastava, R., & Howell, S. H. (2013). Endoplasmic reticulum (ER) stress response and its physiological roles in plants. International journal of molecular sciences, 14(4), 8188–8212.

Dobrev, P.I., Kamínek, M. (2002). Fast and efficient separation of cytokinins from auxin and abscisic acid and their purification using mixed-mode solid-phase extraction. J Chromatogr A, 950:21–9.

Edlund, E., Novak, O., Karady, M., Ljung, K., & Jansson, S. (2017). Contrasting patterns of cytokinins between years in senescing aspen leaves. Plant, cell & environment, 40(5), 622–634.

Epple, P., Mack, A. A., Morris, V. R., & Dangl, J. L. (2003). Antagonistic control of oxidative stress-induced cell death in Arabidopsis by two related, plant-specific zinc finger proteins. Proceedings of the National Academy of Sciences, 100(11), 6831–6836.

Fataftah, N., Bag, P., André, D., Lihavainen, J., Zhang, B., Ingvarsson, P. K., … & Jansson, S. (2021). GIGANTEA influences leaf senescence in trees in two different ways. Plant physiology, 187(4), 2435–2450.

Fataftah, N., Edlund, E., Lihavainen, J., Bag, P., Björkén, L., Näsholm, T., & Jansson, S. (2021). Nitrate fertilization may delay autumn leaf senescence, while amino acid treatments do not. bioRxiv.

Floková, K., Tarkowská, D., Miersch, O., Strnad, M., Wasternack, C., & Novák, O. (2014). UHPLC-MS/MS based target profiling of stress-induced phytohormones. Phytochemistry, 105(July), 147–157.

Fracheboud, Y., Luquez, V., Björkén, L., Sjödin, A., Tuominen, H., & Jansson, S. (2009). The control of autumn senescence in European aspen. Plant physiology, 149(4), 1982–1991.

Gepstein S (2004) Leaf senescence—not just a ‘wear and tear’ phenomenom. Genome Biol 5:212

Großkinsky, D. K., Syaifullah, S. J., & Roitsch, T. (2018). Integration of multi-omics techniques and physiological phenotyping within a holistic phenomics approach to study senescence in model and crop plants. Journal of Experimental Botany, 69(4), 825–844.

Guo, Y., & Gan, S. S. (2012). Convergence and divergence in gene expression profiles induced by leaf senescence and 27 senescence-promoting hormonal, pathological and environmental stress treatments. Plant, Cell & Environment, 35(3), 644–655.

Guo, P., Li, Z., Huang, P., Li, B., Fang, S., Chu, J., & Guo, H. (2017). A tripartite amplification loop involving the transcription factor WRKY75, salicylic acid, and reactive oxygen species accelerates leaf senescence. The Plant Cell, 29(11), 2854–2870.

Heberle, H., Meirelles, G. V., da Silva, F. R., Telles, G. P., & Minghim, R. (2015). InteractiVenn: a web-based tool for the analysis of sets through Venn diagrams. BMC bioinformatics, 16(1), 1–7.

Herrera-Vásquez, A., Salinas, P., & Holuigue, L. (2015). Salicylic acid and reactive oxygen species interplay in the transcriptional control of defense genes expression. Frontiers in plant science, 6, 171.

Howell, S. H. (2013). Endoplasmic reticulum stress responses in plants. Annual review of plant biology, 64, 477–499

Huang, X., Li, Y., Zhang, X., Zuo, J., & Yang, S. (2010). The Arabidopsis LSD1 gene plays an important role in the regulation of low temperature-dependent cell death. New Phytologist, 187(2), 301–312.

Huang, X. X., Zhu, G. Q., Liu, Q., Chen, L., Li, Y. J., & Hou, B. K. (2018). Modulation of plant salicylic acid-associated immune responses via glycosylation of dihydroxybenzoic acids. Plant physiology, 176(4), 3103–3119.

Jansson, S., & Thomas, H. (2008). Senescence: developmental program or timetable?. New Phytologist, 575–579.

Karlberg, A., Englund, M., Petterle, A., Molnar, G., Sjödin, A., Bako, L., & Bhalerao, R. P. (2010). Analysis of global changes in gene expression during activity-dormancy cycle in hybrid aspen apex. Plant Biotechnology, 27(1), 1–16.

Keskitalo, J., Bergquist, G., Gardeström, P., & Jansson, S. (2005). A cellular timetable of autumn senescence. Plant Physiology, 139(4), 1635–1648.

Kopylova, E., Noé, L., & Touzet, H. (2012). SortMeRNA: fast and accurate filtering of ribosomal RNAs in metatranscriptomic data. Bioinformatics, 28(24), 3211–3217.

Lai, Y. S., Renna, L., Yarema, J., Ruberti, C., He, S. Y., & Brandizzi, F. (2018). Salicylic acid-independent role of NPR1 is required for protection from proteotoxic stress in the plant endoplasmic reticulum. Proceedings of the National Academy of Sciences, 115(22), E5203–E5212.

Langfelder, P., & Horvath, S. (2008). WGCNA: an R package for weighted correlation network analysis. BMC bioinformatics, 9(1), 559.

Langfelder, P., Luo, R., Oldham, M. C., and Horvath, S. (2011). Is my network module preserved and reproducible? PLoS Comput. Biol. 7:e1001057. doi: 10.1371/journal.pcbi.1001057

Lee, S., Kim, M. H., Lee, J. H., Jeon, J., Kwak, J. M., & Kim, Y. J. (2020). Glycosyltransferase-Like RSE1 Negatively Regulates Leaf Senescence Through Salicylic Acid Signaling in Arabidopsis. Frontiers in Plant Science, 11, 551.

Li, L., Xing, Y., Chang, D., Fang, S., Cui, B., Li, Q., … & Shen, Y. (2016). CaM/BAG5/Hsc70 signaling complex dynamically regulates leaf senescence. Scientific reports, 6(1), 1–12.

Li J, Brader G, Palva ET (2004) The WRKY70 transcription factor: a node of convergence for jasmonate-mediated and salicylate-mediated signals in plant defense. Plant Cell 16:319– 331

Li, W., Zhang, H., Li, X., Zhang, F., Liu, C., Du, Y., … & Guo, Y. (2017). Intergrative metabolomic and transcriptomic analyses unveil nutrient remobilization events in leaf senescence of tobacco. Scientific reports, 7(1), 1–17.

Li, Z., Zhang, Y., Zou, D., Zhao, Y., Wang, H. L., Zhang, Y., … & Zhang, Z. (2020). LSD 3.0: a comprehensive resource for the leaf senescence research community. Nucleic acids research, 48(D1), D1069-D1075.

Lihavainen, J., Edlund, E., Björkén, L., Bag, P., Robinson, K. M., & Jansson, S. (2020). Stem Girdling Affects The Onset of Autumn Senescence In Aspen In Interaction With Metabolic Signals. Physiologia Plantarum.

Lim, P. O., Kim, H. J., & Gil Nam, H. (2007). Leaf senescence. Annu. Rev. Plant Biol., 58, 115–136.

Love, M. I., Huber, W., & Anders, S. (2014). Moderated estimation of fold change and dispersion for RNA-seq data with DESeq2. Genome biology, 15(12), 1–21.

Lu, H., Gordon, M. I., Amarasinghe, V., & Strauss, S. H. (2020). Extensive transcriptome changes during seasonal leaf senescence in field-grown black cottonwood (Populus trichocarpa Nisqually-1). Scientific reports, 10(1), 1–14.

Luquez, V., Hall, D., Albrectsen, B. R., Karlsson, J., Ingvarsson, P., & Jansson, S. (2008). Natural phenological variation in aspen (Populus tremula): the SwAsp collection. Tree Genetics & Genomes, 4(2), 279–292.

Matías-Hernández, L., Aguilar-Jaramillo, A. E., Marín-González, E., Suárez-López, P., & Pelaz, S. (2014). RAV genes: regulation of floral induction and beyond. Annals of botany, 114(7), 1459–1470.

Michelson, I. H., Ingvarsson, P. K., Robinson, K. M., Edlund, E., Eriksson, M. E., Nilsson, O., & Jansson, S. (2018). Autumn senescence in aspen is not triggered by day length. Physiologia plantarum, 162(1), 123–134.

Morris, K., -Mackerness, S. A. H., Page, T., John, C. F., Murphy, A. M., Carr, J. P., & Buchanan-Wollaston, V. (2000). Salicylic acid has a role in regulating gene expression during leaf senescence. The Plant Journal, 23(5), 677–685.

Nagashima, Y., Iwata, Y., Ashida, M., Mishiba, K. I., & Koizumi, N. (2014). Exogenous salicylic acid activates two signaling arms of the unfolded protein response in Arabidopsis. Plant and Cell Physiology, 55(10), 1772–1778.

Patro, R., Duggal, G., Love, M. I., Irizarry, R. A., & Kingsford, C. (2017). Salmon provides fast and bias-aware quantification of transcript expression. Nature methods, 14(4), 417–419.

Pattison, R. J., & Amtmann, A. (2009). N-glycan production in the endoplasmic reticulum of plants. Trends in plant science, 14(2), 92–99.

Poór, P., Czékus, Z., Tari, I., & Ördög, A. (2019). The multifaceted roles of plant hormone salicylic acid in endoplasmic reticulum stress and unfolded protein response. International journal of molecular sciences, 20(23), 5842.

Poór, P. (2020). Effects of salicylic acid on the metabolism of mitochondrial reactive oxygen species in plants. Biomolecules, 10(2), 341.

Radojicic, A., Li, X., & Zhang, Y. (2018). Salicylic acid: A double-edged sword for programed cell death in plants. Frontiers in plant science, 9, 1133.

Ramirez-Prado, J. S., Abulfaraj, A. A., Rayapuram, N., Benhamed, M., & Hirt, H. (2018). Plant immunity: from signaling to epigenetic control of defense. Trends in plant science, 23(9), 833–844.

Raudvere, U., Kolberg, L., Kuzmin, I., Arak, T., Adler, P., Peterson, H., & Vilo, J. (2019). g: Profiler: a web server for functional enrichment analysis and conversions of gene lists (2019 update). Nucleic acids research, 47(W1), W191-W198.

Ren, K., Feng, L., Sun, S., & Zhuang, X. (2021). Plant mitophagy in comparison to mammals: What is still missing?. International Journal of Molecular Sciences, 22(3), 1236.

Rigault, M., Citerne, S., Masclaux-Daubresse, C., & Dellagi, A. (2021). Salicylic acid is a key player of Arabidopsis autophagy mutant susceptibility to the necrotrophic bacterium Dickeya dadantii. Scientific Reports, 11(1), 1–10.

Rustérucci, C., Aviv, D. H., Holt, B. F., Dangl, J. L., & Parker, J. E. (2001). The disease resistance signaling components EDS1 and PAD4 are essential regulators of the cell death pathway controlled by LSD1 in Arabidopsis. The Plant Cell, 13(10), 2211–2224.

Schäfer, P., & Eichmann, R. (2012). The endoplasmic reticulum in plant immunity and cell death. Frontiers in plant science, 3, 200.

Schurch, N. J., Schofield, P., Gierliński, M., Cole, C., Sherstnev, A., Singh, V., … & Barton, G. J. (2016). How many biological replicates are needed in an RNA-seq experiment and which differential expression tool should you use?. Rna, 22(6), 839–851.

Shannon, P., Markiel, A., Ozier, O., Baliga, N. S., Wang, J. T., Ramage, D., … & Ideker, T. (2003). Cytoscape: a software environment for integrated models of biomolecular interaction networks. Genome research, 13(11), 2498–2504.

Sun, T., Zhang, Y., Li, Y., Zhang, Q., Ding, Y., & Zhang, Y. (2015). ChIP-seq reveals broad roles of SARD1 and CBP60g in regulating plant immunity. Nature communications, 6(1), 1–12.

Sun, T., Zhang, Q., Gao, M., & Zhang, Y. (2014). Regulation of SOBIR 1 accumulation and activation of defense responses in bir1–1 by specific components of ER quality control. The Plant Journal, 77(5), 748–756.

Sundell, D., Mannapperuma, C., Netotea, S., Delhomme, N., Lin, Y. C., Sjödin, A., … & Street, N. R. (2015). The plant genome integrative explorer resource: PlantGenIE. org. New Phytologist, 208(4), 1149–1156.

Svačinová, J., Novák, O., Plačková, L., Lenobel, R., Holík, J., Strnad, M., & Doležal, K. (2012). A new approach for cytokinin isolation from Arabidopsis tissues using miniaturized purification: pipette tip solid-phase extraction. Plant methods, 8(1), 1–14.

Ülker, B., Mukhtar, M. S., & Somssich, I. E. (2007). The WRKY70 transcription factor of Arabidopsis influences both the plant senescence and defense signaling pathways. Planta, 226(1), 125–137.

Valandro, F., Menguer, P. K., Cabreira-Cagliari, C., Margis-Pinheiro, M., & Cagliari, A. (2020). Programmed cell death (PCD) control in plants: New insights from the Arabidopsis thaliana deathosome. Plant Science, 110603.

Vitale, A. (2009). Calreticulins are not all the same. Proceedings of the National Academy of Sciences, 106(32), 13151–13152.

Vogelmann, K., Drechsel, G., Bergler, J., Subert, C., Philippar, K., Soll, J., … & Hoth, S. (2012). Early senescence and cell death in Arabidopsis saul1 mutants involves the PAD4-dependent salicylic acid pathway. Plant physiology, 159(4), 1477–1487.

Wang, J., Ding, J., Tan, B., Robinson, K. M., Michelson, I. H., Johansson, A., … & Ingvarsson, P. K. (2018). A major locus controls local adaptation and adaptive life history variation in a perennial plant. Genome biology, 19(1), 1–17.

Wang, X., Gao, J., Gao, S., Song, Y., Yang, Z., & Kuai, B. (2019). The H3K27me3 demethylase REF6 promotes leaf senescence through directly activating major senescence regulatory and functional genes in Arabidopsis. PLoS genetics, 15(4), e1008068.

Wang, H. L., Zhang, Y., Wang, T., Yang, Q., Yang, Y., Li, Z., … & Li, Z. (2021). An Alternative Splicing Variant of PtRD26 Delays Leaf Senescence by Regulating Multiple NAC Transcription Factors in Populus. The Plant Cell.

Watanabe, M., Tohge, T., Balazadeh, S., Erban, A., Giavalisco, P., Kopka, J., … & Hoefgen, R. (2018). Comprehensive metabolomics studies of plant developmental senescence. In Plant Senescence (pp. 339–358). Humana Press, New York, NY.

Wituszyńska, W., Szechyńska-Hebda, M., Sobczak, M., Rusaczonek, A., Kozłowska-Makulska, A., Witoń, D., & Karpiński, S. (2015). LESION SIMULATING DISEASE 1 and ENHANCED DISEASE SUSCEPTIBILITY 1 differentially regulate UV-C-induced photooxidative stress signalling and programmed cell death in A rabidopsis thaliana. Plant, cell & environment, 38(2), 315–330.

Woo, H. R., Kim, J. H., Kim, J., Kim, J., Lee, U., Song, I. J., … & Lim, P. O. (2010). The RAV1 transcription factor positively regulates leaf senescence in Arabidopsis. Journal of Experimental Botany, 61(14), 3947–3957.

Woo, H. R., Kim, H. J., Nam, H. G., & Lim, P. O. (2013). Plant leaf senescence and death– regulation by multiple layers of control and implications for aging in general. Journal of cell science, 126(21), 4823–4833.

Woo, H. R., Kim, H. J., Lim, P. O., & Nam, H. G. (2019). Leaf senescence: systems and dynamics aspects. Annual Review of Plant Biology, 70, 347–376.

Yin, R., Liu, X., Yu, J., Ji, Y., Liu, J., Cheng, L., & Zhou, J. (2020). Up-regulation of autophagy by low concentration of salicylic acid delays methyl jasmonate-induced leaf senescence. Scientific Reports, 10(1), 1–10.

Yoshioka, K., Kachroo, P., Tsui, F., Sharma, S. B., Shah, J., & Klessig, D. F. (2001). Environmentally sensitive, SA-dependent defense responses in the cpr22 mutant of Arabidopsis. The Plant Journal, 26(4), 447–459.

Zhang, Y., Xu, S., Ding, P., Wang, D., Cheng, Y. T., He, J., Gao, M., Xu, F., Li, Y., Zhu, Z., Li, X., and Zhang, Y. 2010. Control of salicylic acid synthesis and systemic acquired resistance by two members of a plant-specific family of transcription factors. Proc. Natl. Acad. Sci. U.S.A. 107:18220–18225.

Zhang, Z., & Guo, Y. (2018). Hormone treatments in studying leaf senescence. In Plant Senescence (pp. 125–132). Humana Press, New York, NY.

Zhang, Y., Wang, H. L., Li, Z., & Guo, H. (2020). Genetic network between leaf senescence and plant immunity: Crucial regulatory nodes and new insights. Plants, 9(4), 495.

Zhou, M., Lu, Y., Bethke, G., Harrison, B. T., Hatsugai, N., Katagiri, F., & Glazebrook, J. (2018). WRKY70 prevents axenic activation of plant immunity by direct repression of SARD1. new phytologist, 217(2), 700–712.

Zhu, C., Xiaoyu, L., Junlan, G., Yun, X., & Jie, R. (2020). Integrating transcriptomic and metabolomic analysis of hormone pathways in Acer rubrum during developmental leaf senescence. BMC Plant Biology, 20(1), 1–22.

